# RISOP, a Reference-Assisted Approach for Enhanced Identification of Oxidized Phospholipids

**DOI:** 10.64898/2026.06.01.728773

**Authors:** Zixing Chen, Andrew Erickson, Natalie Ito, Xiaoai Zhao

## Abstract

Oxidized phospholipids (OxPLs) play critical roles in inflammation, ferroptosis, and other oxidative stress-associated processes, yet their systematic characterization in biological systems remains a major analytical challenge owing to their low abundance and vast structural diversity. Here we report RISOP (Reference-Assisted Identification of Sample-specific Oxidized Phospholipids), an untargeted LC-MS/MS workflow that leverages chemically enriched OxPL reference pools to substantially improve OxPL annotation. Reference pools encompassing diverse oxidative modifications and a wide abundance range were experimentally generated using Fenton reaction and H_2_O_2_ treatment, providing broad coverage of OxPLs. Reference-assisted integrative analysis yielded a more than twofold increase in identified OxPL species in biological samples under elevated oxidative stress, as demonstrated in ML210-treated cells, a model of ferroptotic stress. We further show that the widely used BODIPY C11 lipid peroxidation probe captures cellular oxidative burden in only a subset of OxPLs identified by RISOP, highlighting the importance of untargeted, comprehensive OxPL profiling. Overall, RISOP provides a versatile platform readily applicable to other classes of oxidized complex lipids for comprehensive characterization in physiological and pathological contexts.

**Abstract Graphic:** 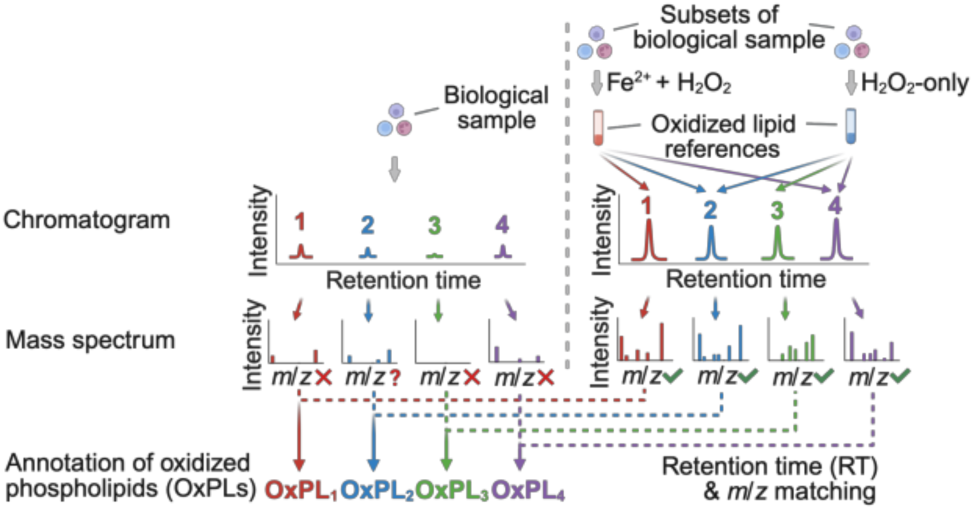

## INTRODUCTION

Oxidized lipids are a structurally diverse group of molecules produced by chemical or enzymatic oxidation of lipids.^1^ They can be broadly classified into oxidized free fatty acids (oxylipins) and oxidized complex lipids, including oxidized phospholipids (OxPLs) and glycerol lipids.^1–3^ Among oxidized complex lipids, OxPLs constitute one of the most structurally diverse classes of oxidized lipids in biological systems.^4^ OxPLs can be formed through lipid peroxidation, a free radical chain reaction consisting of initiation, propagation, and termination steps.^5, 6^ During initiation, reactive radicals abstract hydrogen atoms from the fatty acyl chains of phospholipids to form lipid radicals (L•). These lipid radicals can react with oxygen to generate lipid peroxyl radicals (LOO•), which propagate the reaction by abstracting hydrogen atoms from neighboring lipids, producing lipid hydroperoxides (LOOH) and new lipid radicals. The resulting LOOH can undergo rearrangement, truncation, or further modification, generating a variety of OxPL species with hydroxy (<OH>), hydroperoxy (<OOH>), carboxy (<COOH>) or keto (<oxo>) modifications.^5, 6^

To evaluate lipid peroxidation in biological systems, several cellular assays have been developed that detect intermediates formed during lipid peroxidation. For example, BODIPY C11 reacts with lipid radicals,^7^ whereas Liperfluo detects lipid hydroperoxides (LOOH).^8^ In addition, secondary breakdown products from LOOH such as 4-hydroxynonenal (4-HNE) and malondialdehyde (MDA) can be detected by antibody-based assays.^9, 10^ However, these methods only provide proxy measures of overall lipid peroxidation that do not distinguish the contribution from each pool of oxidized lipids (e.g., oxylipins vs. OxPLs). Furthermore, these assays do not provide identification of the oxidized lipids themselves, missing key information about lipid class, the exact side chain(s) that were oxidized, and the types of oxidative modifications, all of which are critical in understanding the diverse functional roles of OxPLs in biological contexts.

OxPLs have been implicated in biological processes including inflammation,^6, 11, 12^ autophagy,^13^ and cell death pathways^14–16^ that are closely associated with cardiovascular diseases,^17–19^ neurodegenerative diseases^20, 21^ and aging.^22, 23^ Mass spectrometry-based approaches remain the gold standard in analyzing OxPLs.^24^ However, the extremely low abundance and extensive chemical diversity pose significant challenges for detection and identification of OxPLs. Moreover, while robust targeted lipidomics approaches have been widely used in the analytical studies of oxylipins,^25, 26^ a similar workflow cannot be readily adopted for OxPL analysis. This is primarily due to the greater structural complexity of OxPLs (>1000 predicted species)^27–29^ compared to oxylipins (∼100),^30^ compounded by the lack of authentic chemical standards for most OxPLs.^31^ Although mass spectrometry-based methods for OxPL analysis have been reported,^32–36^ these approaches are limited by some forms of narrow lipid class coverage,^32, 36^ workflow or instrumentation complexity,^32, 34, 35^ and applicability restricted to cultured cell systems.^33^ Collectively, these limitations highlight a critical unmet need in developing a streamlined, untargeted lipidomics strategy capable of comprehensive, high-coverage OxPL detection and identification – one that can ultimately advance our mechanistic understanding of OxPL biology.

In this study, we developed RISOP (reference-assisted identification of sample-specific oxidized phospholipids). We established a platform for comprehensive and enhanced identification of OxPLs in an untargeted lipidomics workflow. RISOP integrates two complementary components. First, chemically enriched OxPL reference pools that are generated from the same biological matrix using two oxidation reactions provide high-quality tandem mass spectra and retention-time anchors for OxPLs that are otherwise too scarce to be annotated directly. Second, an *in silico* spectral library, predicted from the non-oxidized lipids of each sample, constrains annotation to biologically plausible species and minimizes false annotation. In addition, RISOP does not require modification of instrumentation or separate setup for data acquisition. Moreover, the reference-generation strategy gives RISOP broad applicability and allows it to be extended beyond cultured cell systems. Overall, this approach expands analytical capability in studies of OxPLs, and enables functional investigations of OxPLs in physiology and disease.

## MATERIALS AND METHODS

### Cell Culture and Preparation for Lipidomics

KP4 cell (passage 15–20) was a generous gift from Dr. Mandar Muzumdar (Yale University). Cells were cultured in 6-well plates in high-glucose Dulbecco’s modified Eagle’s medium (Gibco, 11965-092), supplemented with 10% fetal bovine serum (Gibco, A56708-81) and 1% penicillin streptomycin (Gibco, 15140-122), and maintained in an incubator containing 5% CO_2_ at 37 °C. KP4 cells at 70% confluence were washed twice with 2 mL ice-cold phosphate-buffered saline (PBS, Gibco, 10010-023) and scraped into 400 μL of PBS, immediately flash-frozen on dry ice, and stored at −80 °C until use.

### Preparation of Oxidized Lipid References

Fenton reactions were conducted following a previously reported protocol with some modifications.^37^ First, 400 μL of cell suspension in PBS from one well of the 6-well plates was thawed on ice. Next, 25 μL of 10 M H_2_O_2_ (Thermo Fisher Scientific, L14000.AP) and 75 μL of 333.33 mM of FeCl_2_ (Sigma, 44939-50G) were added to cell suspension for a final concentration of 500 mM of H_2_O_2_ and 50 mM of FeCl_2_. For H_2_O_2_-only reactions, the same amounts of thawed cell suspension in PBS and H_2_O_2_ were used, and an additional 75 μL of PBS was used to substitute for FeCl_2_ solution. Samples were subsequently incubated at 37 °C in the dark for 0 min, 1 min, 20 min, 4 h, 24 h, and 72 h. Following incubation, samples were immediately processed for lipid extraction.

### Lipid Extraction

Lipids were extracted using a 2-phase, liquid-liquid extraction system – a modified Folch method.^38^ All samples were randomized prior to extraction. All chemicals used for extraction were LC/MS grade unless otherwise indicated. First, 300 μL of methanol (Fisher chemical, A456-4) containing 0.5 μL of deuterated internal standards (EquiSPLASH® mix, Avanti Polar Lipids, 330731), 0.5 μL of cardiolipin standard (Avanti Polar Lipids, 791108C), 0.5 μL of oleic acid standard (Cayman chemical, 9000432), and 0.1 mg/mL 2,6-Di-tert-butyl-4-methylphenol (BHT, Thermo Fisher Scientific, 112992500) were added to 400 μL of cell suspension in PBS. The mixture was sonicated for 30 sec in a water bath ultrasonicator (VWR, 97042-960) and then rested on ice for 30 sec, repeated three times. Then 600 μL of ice-cold chloroform (VWR, BDH83627.400) was added to each sample. The resulting mixture was vortexed at 4 °C for 30 min. Phase separation was subsequently achieved by centrifugation at 3,000 RPM for 10 min at 4 °C. The lower organic phase was collected and dried under a stream of nitrogen. Dried lipids were then reconstituted in 150 μL of a 9:1 (v/v) methanol/toluene (Fisher chemical, T290-1) solution for liquid chromatography–tandem mass spectrometry (LC-MS/MS) analysis.

### LC-MS/MS Lipidomics

Lipidomic analysis was conducted in accordance with guidelines of the Lipidomics Standards Initiative (LSI) (https://lipidomicstandards.org/), and all lipidomics-related information is included in the accompanying LSI reporting checklist^39^ (Table S1). LC-MS/MS analysis was performed on a Vanquish UPLC system coupled to an Orbitrap Exploris 120 mass spectrometer (Thermo Fisher Scientific). Samples were randomized prior to analysis. Lipid separation was achieved on a Hypersil GOLD C18 HPLC column (100 mm × 1 mm, 1.9 μm, Thermo Fisher Scientific, 25002-101030) with a flow rate of 120 μL/min. The column temperature was maintained at 45 °C, and the injection volume was 1.5 μL. Mobile phase A is composed of 60:40 (v/v) acetonitrile (Fisher chemical, A955-4)/water (Fisher chemical, W6-4), and mobile phase B is composed of 88:10:2 (v/v/v) isopropanol (Fisher chemical, A461-4)/acetonitrile/water. Both mobile phases were supplemented with 0.1% formic acid (Fisher chemical, A117-50) and 7.5 mM ammonium acetate (Fisher chemical, 198759). The chromatography gradient was as follows: 0–3 min, 10% B; 3–5 min, 10–43% B; 5–5.1 min, 43–55% B; 5.1–15.1 min, 55–65% B; 15.1–21.1 min, 65–85% B; 21.1–23.1 min, 85–100% B; 23.1–28.1 min, 100% B; 28.1–28.2 min, 100–10% B; 28.2–31.2 min, 10% B. The expected LC peak width was set to 8 sec. Mass spectra were acquired in both positive and negative ion modes in two different runs using a heated electrospray ionization (HESI) source. The spray voltages were set to 3,500 V and 2,500 V for positive and negative modes, respectively. The sheath, auxiliary, and sweep gas flow rates were set to 35, 7, and 0 (arbitrary units), respectively. MS_1_ full scans and MS_2_ tandem mass spectra were acquired in all samples. For MS_1_ full scans, the resolution was set to 120,000, with a scan range of 200–1700 *m/z* and an RF lens of 70%. The microscan number was set to 1, with the automatic gain control (AGC) target set to “Standard” and the maximum injection time set to “Auto.” Data were acquired in profile mode. Precursor ions with intensities greater than 2.0 × 10⁴ were selected for fragmentation using Top 4 data-dependent acquisition. MS_2_ scans were acquired at a resolution of 30,000 with the auto-extended scan range mode enabled. The isolation window was set at 1.2 *m/z*. Stepwise normalized collision energies of 20%, 30%, and 40% were used for fragmentation. The microscan number was set at 1. The automatic gain control (AGC) target was set to “Standard,” and the maximum injection time was set to “Auto.” Data were acquired in profile mode. A dynamic exclusion of 8.5 s was applied to prevent repeated fragmentation of the same precursor ion. For quality control, a pooled sample was repeatedly injected after every 10 samples throughout the analytical runs, and the coefficient of variation (CV) of internal standard intensities was used to assess the stability of the LC-MS system. Calculated CVs of all internal standards in this study were <6%. Retention time drift was monitored by the retention time of internal standards. Calculated retention time drift was between 0.01–0.03 minutes.

### Identification of Non-Oxidized Lipids Using MS-DIAL

Lipidomics data files were processed using MS-DIAL software (version 5.5.250221) for the identification of non-oxidized lipids. The analysis was conducted within the software’s lipidomics framework. Centroid data type was selected for both MS_1_ and MS_2_. The MS_1_ and MS_2_ tolerance were set at 0.01 and 0.025 Da, respectively. The minimum peak height was set to 1000 amplitude, and the mass slice width was set to 0.05 Da. Linear weighted moving average was used as the smoothing method, with a smoothing level of 3 scans and minimum peak width of 5 scans. The non-oxidized lipids were identified using the built-in default spectral library (Msp20250218112233_NCDK_conventional_converted_dev_1) in MS-DIAL. For lipid identification, thresholds for dot product score were set at 150, weighted dot product score was set at 150, reverse dot product score was set at 500, matched spectrum percentage was set at 0%, and minimum number of matched spectra was set at 1. HCOONH4 was selected as the solvent type for searching, as we observed that phospholipids with [M+HCOO]^-^ adducts have higher intensities than those with [M+CH_3_COO]^-^ adducts during ionization, likely due to the presence of formic acid in mobile phases.^40, 41^ For positive mode, [M+H]^+^, [M+NH_4_]^+^, [M+Na]^+^, and [M+H-H_2_O]^+^ adducts were selected for annotation. For negative mode, the [M-H]^-^, [M+HCOO]^-^, and [M+CH_3_COO]^-^ adducts were selected for annotation. Samples were aligned to the pooled sample, with a retention time tolerance of 0.1 min and MS_1_ tolerance of 0.015 Da. To minimize misannotation, all identified lipids were manually validated by inspecting the tandem mass spectra for fragment ions diagnostic of the assigned headgroup and fatty acyl chains (in the accompanying LSI reporting checklist [Table S1] and Table S2). All non-oxidized lipids were reported at the molecular species level (see “*Lipidomics Nomenclature*” below).

### Construction of Sample-specific OxPL Spectral Library

Sample-specific OxPL spectral library was constructed from identified lipids of the following classes: phosphatidylcholine (PC), ether-linked phosphatidylcholine (PC-O), phosphatidylethanolamine (PE), ether-linked phosphatidylethanolamine (EtherPE-O), ether-linked phosphatidylethanolamine (EtherPE-P), phosphatidylglycerol (PG), phosphatidylinositol (PI), phosphatidylserine (PS), lysophosphatidylcholine (LPC), ether-linked lysophosphatidylcholine (EtherLPC-O), lysophosphatidylethanolamine (LPE), and ether-linked lysophosphatidylethanolamine (EtherLPE-O). The list of identified non-oxidized lipids with their lipid classes was imported into LPPtiger 2 for *in silico* oxidation, with a maximum number of modification sites set at 3, and the maximum number of O, OH, oxo, OOH, and epoxy modifications set at 4, 2, 1, 1, and 0, respectively. The resulting theoretical tandem mass spectral library, exported from LPPtiger 2 in JSON format, was organized to include the diagnostic ions corresponding to oxidized lipid precursor ion, headgroup fragment, unoxidized fatty acyl chain(s), and oxidized fatty acyl chain(s) of the OxPLs, and converted to MSP format using a custom script in R (version 4.5.0). For example, for PC(16:0_16:1<OH>), the following diagnostic ions are included: *m/z* 224.0682-headgroup, *m/z* 255.2330-unoxidized side chain (16:0), *m/z* 269.2122-oxidized side chain (16:1<OH>), *m/z* 251.2017-oxidized side chain with H_2_O loss (16:1<OH> − H_2_O), *m/z* 792.5396-precursor (PC(16:0_16:1<OH>) with [M+HCOO]^-^ adduct), and *m/z* 732.5185-precursor with neutral loss of CH_3_COOH.

### Identification of OxPLs in Fenton and H_2_O_2_-only Oxidized Lipid References

OxPL identification in separate Fenton and H_2_O_2_-only references: Lipidomics data files obtained from Fenton or H_2_O_2_-only reactions were analyzed separately in MS-DIAL. Parameters used were identical to those described above for non-oxidized lipids, except that the sample-specific OxPL spectral library constructed above was used in place of the default spectral library. All identified oxidized lipids were manually validated by inspecting the tandem mass spectra for fragment ions diagnostic of the assigned headgroup and fatty acyl chain(s) (Table S1 and Table S2). OxPLs were reported at the molecular species level when both fatty acyl chains were identified in the spectra and at the species level when only one fatty acyl chain was identified.

Only OxPLs that were detected in at least three samples across different time points of each reference type were kept. To minimize false annotation of background noise, coefficient of variation (CV)-based filtering was applied separately to non-oxidized lipids and OxPLs. Non-oxidized lipids were retained if they had a CV <20% for at least one treatment time point (including 0 h) in either Fenton or H_2_O_2_-only references. OxPLs were retained if they had a CV <20% for at least one treatment time point (excluding 0 h) in either reference set. Calculation of CV values is described below in “*Quantification of Non-oxidized and Oxidized Lipids in Fenton Reaction and H_2_O_2_-only Oxidized Lipid References*”. Identified lipids from the 2 separate analyses after the removal of potential in-source fragmentation (ISF) products (see “*Hydrogen-based Kendrick Mass Defect (H-KMD) Analysis*” below) were used to generate the Venn diagram in Figure 2B.

OxPL identification by combined analysis of Fenton and H_2_O_2_-only references: Lipidomics data files from the Fenton reactions, H_2_O_2_-only reactions, and ML210-treated samples (see “*ML210 treatment*” below) were analyzed together in MS-DIAL for integrative analysis and identification. This approach was used to ensure consistent OxPL annotation based on retention time and *m/z* matching across biological samples and oxidized lipid references. The MS-DIAL settings and lipid validation steps were identical to those described above for OxPL identification. This workflow provides a list of OxPLs detected in Fenton reaction and H_2_O_2_-only reference samples. Only OxPLs that were detected in at least three samples across different time points of each reference type were kept. Similarly, CV-based filtering was applied separately to non-oxidized lipids and OxPLs. Non-oxidized lipids were retained if they had a CV <20% for at least one treatment time point (including 0 h) in either Fenton or H_2_O_2_-only references. OxPLs were retained if they had a CV <20% for at least one treatment time point (excluding 0 h) in either reference set. The resulting filtered OxPLs from the Fenton and H_2_O_2_-only references after the removal of potential ISF products were used to generate the Venn diagram in Figure 3A.

### Hydrogen-based Kendrick Mass Defect (H-KMD) Analysis

H-KMD analysis was performed to visualize the differences of retention time and H-KMD between oxidized phospholipids and their corresponding non-oxidized precursors. For each lipid, the observed *m/z* was first multiplied by the nominal-to-exact hydrogen mass ratio (1.000000/1.007825032) to calculate the Kendrick mass, which was subsequently rounded to the nearest integer to obtain the nominal Kendrick mass. H-KMD was then calculated as the difference between the Kendrick mass and the nominal Kendrick mass. The resulting H-KMD values of the OxPLs were plotted against their retention times. OxPLs were annotated as potential ISF products if they met any of the following criteria: (1) they had the same (within a ±0.05 min tolerance) or a later retention time than the corresponding non-oxidized lipids; (2) they had the same retention time (within a ±0.05 min tolerance), as lipids with the same backbone but more labile oxidative modifications; or (3) their retention times were significantly later (>2 min) than the corresponding isomers with the same modifications.

### Quantification of Non-oxidized and Oxidized Lipids in Fenton Reaction and H_2_O_2_-only Oxidized Lipid References

We observed that peak areas of deuterated internal standards included in the Fenton- and H_2_O_2_-only samples showed large variation between replicates, likely due to degradation in the presence of H_2_O_2_ and/or Fe^2+^. Therefore, we used relative quantification by normalizing each lipid to the total peak area of all detected lipids in analyzing oxidized and non-oxidized lipids from these samples. Peak integration of all OxPLs was manually inspected. When a peak was only partially integrated by MS-DIAL, the peak start and end points were manually adjusted to cover the entire chromatographic peak, using samples with clearly defined peak boundaries as references.

For cardiolipin (CL), PE, LPE, PG, PI, PS, FA, EtherPE-O, oxidized PE (OxPE), oxidized PG (OxPG), and oxidized PS (OxPS), quantification was done using [M−H]⁻ adduct. For PC, LPC, EtherPC-O, EtherPE-P, EtherLPC-O, and EtherLPE-O, quantification was done using [M+H]⁺ adduct. For DG and TG, quantification was done using [M+NH4]⁺ adduct. For OxPC with oxidative modifications other than carboxy (<COOH>) modification and oxidized EtherPC (OxEtherPC), quantification was done using [M+HCOO]⁻ adducts rather than [M+H]⁺ adducts (as used in their non-oxidized counterparts) because the detected lipid intensity is higher in negative mode owing to oxidation modifications.^42^ For oxidized PC (OxPC) with carboxy (<COOH>) modifications, quantification was done using [M−H]⁻ adducts, because the carboxyl group provides an acidic site that facilitates deprotonation in negative-ion mode.^43, 44^ CV values were calculated separately for each lipid at each treatment time point using normalized intensities across replicates.

### Lipidomics Nomenclature

Lipids were annotated according to the nomenclature of LIPID MAPS (https://www.lipidmaps.org/). At the molecular species level, lipids were reported with their constituent fatty acyl chains, with carbon number and double bond number separated by a colon (e.g., PC(16:0_18:1)). At the species level, lipids were reported by the summed carbon number and double bond number of all acyl chains (e.g., PC(34:1 <oxo>)). For phospholipids, the fatty acid side chains were separated by an underscore (“_”), indicating that the sn-position of side chains were not specified. Using our mass spectrometry setup, alkyl-linked LPE and LPC species (e.g. LPE(O-18:1)) exhibited the same tandem mass spectra as alkenyl-linked LPE and LPC species with 1 less double bond (e.g. LPE(P-18:0), as previously reported.^45^ For those species, both annotations are listed and separated by “or” (e.g. LPE(O-18:1) or LPE(P-18:0)).

Oxidative modifications were indicated in angle brackets (“<>”), where OH, oxo, OOH, and COOH denote hydroxy, keto, hydroperoxy, and carboxy modifications, respectively. Isomeric species with identical chemical formula but distinct retention times were distinguished by assigning alphabetical suffixes (e.g., (a), (b), (c)) in the order of increasing retention time.

### Lipidomics Reporting

The lipidomic data were uploaded to Metabolomics Workbench (Study ID ST004800; DataTrack ID 7357) and can be accessed at: http://dx.doi.org/10.21228/M8NP12

### Ferroptosis-induced Cytotoxicity Assay

KP4 cells were seeded onto 96-well plates at approximately 5000 cells per well. 24 hours later, cells were treated with ML210 (Sigma, SML0521) dissolved in dimethyl sulfoxide (DMSO) at different concentrations as indicated and incubated for 24 hours. For rescue assay, 5 μM ferrostatin-1 (Sigma, SML0583) solution in DMSO was co-administered with different concentrations of ML210 for 24 hours. Following the treatment, 10 μL of CCK-8 solution (Cell Counting Kit 8, abcam, ab228554) was added to each well and the absorbance at 460 nm was measured using a Varioskan LUX multimode microplate reader (Thermo Fisher Scientific) after 1 hour of incubation at 37 °C.

### ML210 Treatment

KP4 cells were cultured in 6-well plates. When reaching 70% confluence, cells were treated with 10 μM ML210 in culture medium for 0, 20, 45, 80, 120, or 270 minutes. Following treatment, medium was aspirated, and cells were washed with 2 mL ice-cold PBS twice. For untargeted lipidomic analysis, one well of cells was collected for each biological replicate, with three biological replicates prepared for each time point, and analyzed by LC-MS/MS following the protocols described above.

### Cell Viability Assay by Flow Cytometry

The viability of cells treated with 10 μM ML210 in culture medium for 0, 20, 45, 80, 120, or 270 minutes was assessed using flow cytometry. Debris was first excluded based on forward and side scatter characteristics, followed by removal of doublets using singlet gating. Live cells were defined as DAPI-negative events, and cell viability was calculated as the percentage of DAPI-negative cells within the singlet population.

### Identification of Non-oxidized and Oxidized Lipids in ML210-treated Samples

Non-oxidized lipids from ML210-treated samples were identified following the same protocols as described above. For OxPL identification, lipidomics data files from ML210-treated samples were analyzed in MS-DIAL either alone or together with lipidomics data files from oxidized lipid reference samples through four independent analyses: 1) ML210-treated samples alone; 2) ML210-treated samples combined with Fenton reference samples; 3) ML210-treated samples combined with H_2_O_2_-only reference samples; and 4) ML210-treated samples combined with both Fenton and H_2_O_2_-only reference samples. The MS-DIAL settings and lipid validation procedures were identical to those described above for OxPL identification (in “*Identification of OxPLs in Fenton and H_2_O_2_-only Oxidized Lipid References*” section). CV-filtering was applied to minimize false annotation of background noise. Non-oxidized lipids were retained if they had a CV <20% for at least one treatment time point (including 0 min). OxPLs were retained if they had a CV <20% for at least one treatment time point (excluding 0 min). The calculation of CV values is described below in “*Quantification of Non-oxidized and Oxidized Lipids in ML210-treated cells*”. Identified lipids after the removal of potential ISF products (see “*Hydrogen-based Kendrick Mass Defect (H-KMD) Analysis*” above) were used to generate the stacked bar plot for comparison of number of identified OxPLs.

### Protein Quantification in ML210-treated Samples

Aqueous phase was removed following Folch extraction. The remaining protein interphase was washed once with 400 μL ice-cold methanol at room temperature, then centrifuged at 3000 RPM for 10 min at 4 °C and dried down under nitrogen stream, before dissolving in 100 μL of 5% SDS (Thermo Fisher Scientific, J63394.AK). Protein content was measured using a Pierce BCA assay kit (Thermo Fisher Scientific, 23225) as previously reported following vendor’s instructions.^46^

### Quantification of Non-oxidized and Oxidized Lipids in ML210-treated cells

Non-oxidized endogenous lipids were reported in molar concentrations based on the intensity ratio of the same ion adduct between endogenous lipids and the deuterated lipid standard within the same lipid class with known concentration. The lipid adducts used for quantification of non-oxidized lipids were the same as those used in Fenton and H_2_O_2_-only references. Furthermore, lipid molar concentrations were then normalized to cellular protein content and expressed as nmol/mg protein to account for variation in sample preparation. CV values were calculated separately for each lipid at each treatment time point using normalized molar concentrations.

Peak integration of all OxPLs was manually inspected. Manual peak integration was performed as described above (in “*Quantification of Non-oxidized and Oxidized Lipids in Fenton reaction and H_2_O_2_-only Oxidized Lipid References*”). Molecular concentrations of oxidized lipids were calculated based on the intensity ratio of the same ion adduct between oxidized lipids and the deuterated lipid standard within the same lipid class with known concentration. The lipid adducts used for quantification of OxPLs were the same as those used in Fenton and H_2_O_2_-only references. CV-filtering was applied to minimize false annotation of background noise. CV of normalized molar concentrations was calculated across replicates at each treatment time point (excluding 0 h). OxPLs were retained if their CV was below 20% for at least one of the time points.

### Principal Component Analysis (PCA) of Lipidomic Datasets

PCA was performed using the prcomp() function in R (version 4.5.0) on log2-transformed data of normalized peak area for oxidized reference samples, and log2-transformed data of normalized lipid molar concentration for ML210-treated cells. All PCA plots were made to visualize samples on principal component 1 (PC1) and principal component 2 (PC2).

### K-means Clustering on OxPLs

K-means clustering was performed on Z scores of OxPL abundance using the kmeans() function in R (version 4.5.0). Specifically, Z score, the number of standard deviations a data point is away from the mean, was calculated for each OxPL across all replicates and time points, and the mean Z score at each time point was used for clustering. For oxidized reference samples, total abundances of normalized peak area of commonly detected 57 OxPLs from Fenton and H_2_O_2_-only references were used as input. K-means clustering was then performed on the Z score-transformed data with the number of clusters set to 9. For ML210-treated cells, normalized lipid molar concentrations of 22 identified OxPLs were used as input. K-means clustering was then performed on the Z score-transformed data with the number of clusters set to 3.

### Heatmap Visualization

Heatmap visualization was performed using ComplexHeatmap package (version 2.24.1). For Fenton and H_2_O_2_ references, the intensity of each OxPL was normalized to the total OxPL intensity within the same sample to calculate its relative abundance. In each replicate, relative abundance of OxPLs was averaged across treatment time points (excluding 0 h). Missing values were imputed with 5% of the minimum non-missing value across all replicates to enable row-wise Z score transformation. The resulting Z scores were used for heatmap visualization, with original missing values indicated as not detected rather than displayed based on imputed values. OxPLs were grouped according to oxidative modification type, and columns were grouped by reference type.

For ML210-treated cells, molar concentration of each OxPL was transformed into Z score across samples for visualization. Rows were grouped based on the K-means clusters determined above. Columns were arranged based on ML210 treatment time points in increasing order.

### Lipid Peroxidation Assay

For BODIPY C11 staining, cells were incubated with medium containing 5 μM BODIPY 581/591 C11 (Thermo Fisher Scientific, D3861) in DMSO for 30 min at 37 °C. Cells were then trypsinized by 0.25% trypsin-EDTA (Gibco, 25200-056), washed with 2 mL of Hank’s balanced salt solution (Gibco, 14175-095), and analyzed using a BD Symphony flow cytometer. At least 100,000 events were analyzed for each sample. Reduced and oxidized BODIPY 581/591 C11 signal was measured in PE-Texas Red channel (610/20 nm) and FITC channel (515/25 nm), respectively. Peroxidation-positive cells were defined as the proportion of FITC-positive population of total live cells based on gating determined from untreated controls.

### Correlation between OxPL Abundance by LC-MS/MS and Peroxidative Burden by BODIPY C11 Peroxidation Assay in ML210-treated Cells

Pearson correlation analysis was performed between percentage of peroxidation-positive cells and total OxPL abundance. Specifically, total OxPL concentration in each sample of individual time point was first calculated by aggregating normalized molar concentrations of all identified

OxPLs. Next, Z scores of total OxPL abundance were then calculated and averaged within each time point. Similarly, Z scores of peroxidation-positive cells treated with ML210 were calculated and averaged within each time point. Subsequently, the mean Z scores of peroxidation-positive cells were compared to the mean Z scores of total OxPL abundance by Pearson correlation.

Pearson correlation was performed between percentage of peroxidation-positive cells and each of the 3 OxPL clusters identified by K-means clustering. For each cluster, total concentration of OxPLs was first calculated by aggregating normalized molar concentrations of all identified OxPLs. Next, Z scores of total OxPL abundance were then calculated and averaged within each time point. Lastly, the mean Z scores of peroxidation-positive cells across ML210 treatment times (described above) were compared to the mean Z scores of total OxPL abundance of each OxPL cluster by Pearson correlation.

### Statistical Analyses

For comparisons between two groups, *P* values were calculated using Welch’s *t*-test following confirmation of data normality. For comparisons involving three or more groups, *P* values were calculated using Welch’s *t*-test following confirmation of data normality and adjusted for multiple comparisons using the Benjamini–Hochberg method.

## RESULTS AND DISCUSSION

### Reference-assisted Identification of Sample-specific Oxidized Phospholipid (RISOP)

In biological samples, non-oxidized phospholipids account for approximately 40–70% of the total mammalian lipidome.^47, 48^ In contrast, oxidized phospholipids (OxPLs) are very low in abundance, generally estimated to be less than 5% of their non-oxidized counterparts.^49–51^ Reliable detection and accurate annotation have long been a challenge in the field, as these low-abundance lipids often fail to generate sufficiently informative tandem mass spectra (MS/MS) that are necessary for structural annotation (**Figure 1A**). To overcome this challenge, we reasoned that creating a pool of oxidized phospholipid references would yield high-quality spectral data. This reference pool was designed to enrich OxPL species present in greater number and abundance, which is critical for the identification of otherwise difficult-to-detect OxPLs from biological samples. We generated OxPL references by subjecting a subset of biological samples to two chemical reactions – Fenton reaction, which is initiated by Fe^2+^ and H_2_O_2_,^37, 52^ as well as treatment with H_2_O_2_ alone. We used these two reactions because it has been reported that Fenton reaction induces strong and non-selective oxidation by producing highly reactive hydroxyl radicals via ferrous ion-mediated reactions,^53–55^ whereas oxidation by H_2_O_2_ alone (H_2_O_2_-only hereafter) proceeds under metal-limited conditions and is therefore comparatively milder, likely relying on endogenous metal ions present in biological samples.^56–58^ Therefore, chemically-enriched OxPL reference pools from these two reactions will likely expand the reference repertoire and enhance our ability to identify and annotate OxPLs.

**Figure 1.**
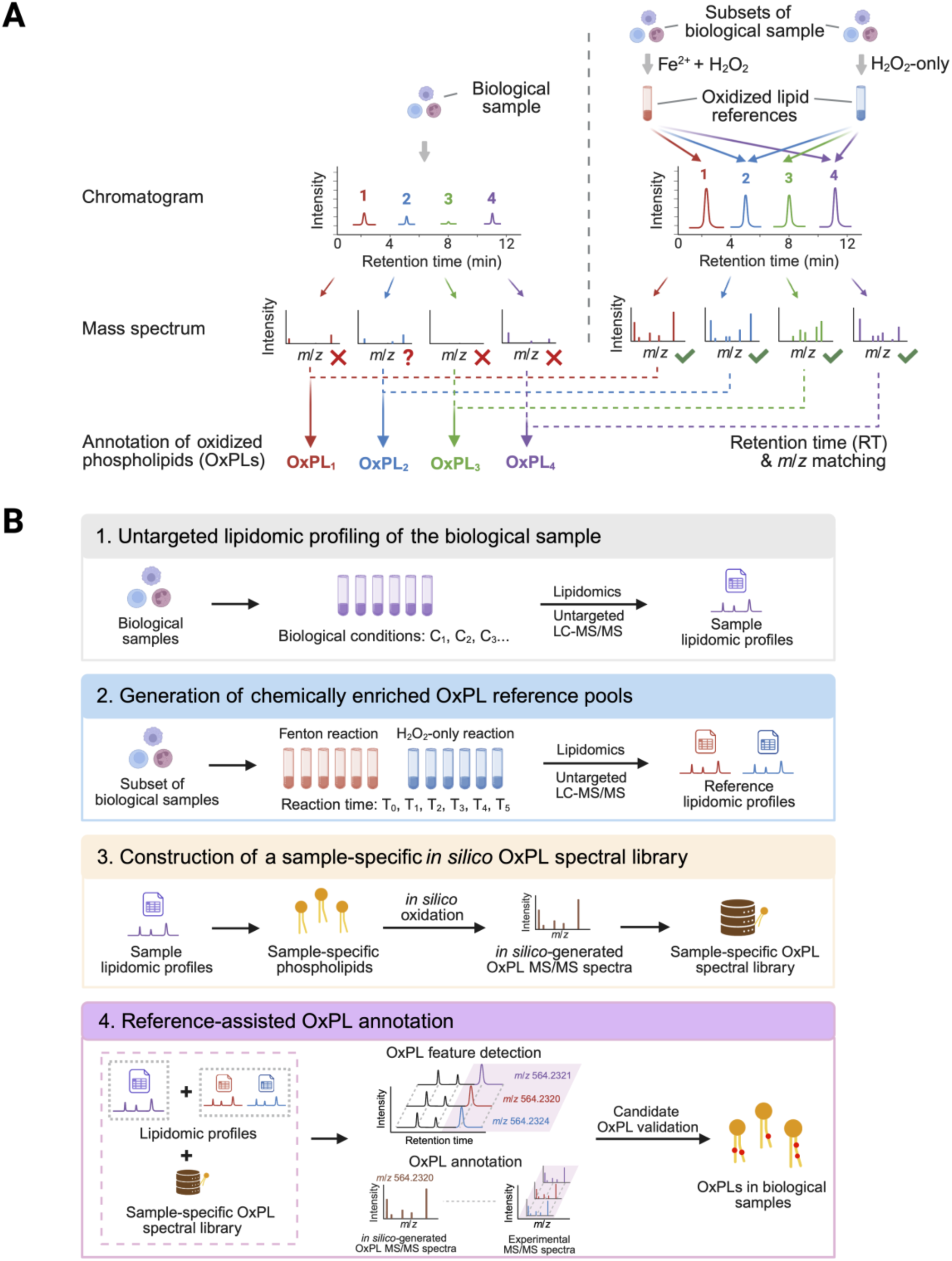
Reference-assisted identification of sample-specific oxidized phospholipid (RISOP). (**A**) In biological samples, oxidized phospholipids (OxPLs) are often present at low abundance. Consequently, their MS spectra are often insufficiently informative, making their annotation challenging (left panel). To overcome this challenge, we experimentally generate a reference pool of oxidized lipids by 2 oxidation reactions. These references are chemically enriched for oxidized phospholipid (OxPL) species in greater number and abundance, enabling identification of low-abundance OxPLs in biological samples through retention time (RT) and mass-to-charge ratio (*m/z*) matching to the oxidized lipid references (right panel). (**B**) Workflow of RISOP. 1) Untargeted lipidomic profiling of the biological sample. Lipidomic profiles of biological samples were obtained by untargeted LC-MS/MS lipidomics. 2) Generation of chemically enriched OxPL reference pools. Oxidized lipid references were generated by subjecting a subset of the biological samples to Fenton reaction or H_2_O_2_ treatment alone (H_2_O_2_-only) for different reaction times, followed by LC-MS/MS lipidomic analysis. 3) Construction of a sample-specific *in silico* OxPL spectral library. The endogenous non-oxidized phospholipids identified from the biological samples from step 1 were used for *in silico* oxidation. The *in silico*-generated tandem mass spectra containing diagnostic ions of each OxPL species were used to generate a sample-specific oxidized phospholipid spectral library. 4) Reference-assisted OxPL annotation. OxPL features were first detected by retention time and *m/z* matching between biological samples and the experimental reference pools. These features were then annotated by matching their experimental MS/MS spectra against *in silico* MS/MS spectra generated from the sample-specific spectral library. OxPLs identified in biological samples were reported following spectral validation to remove misannotated OxPLs.

We refer to this method as Reference-Assisted Identification of Sample-specific Oxidized Phospholipids (RISOP). RISOP integrates four steps (**Figure 1B**), combining empirical reference spectra with an *in silico*–defined search space to enable annotation of low-abundance OxPLs that would otherwise lack interpretable MS/MS evidence. **1) Untargeted lipidomic profiling of the biological sample.** Lipids are extracted and profiled by LC-MS/MS; non-oxidized lipids are annotated against established libraries and used to define the sample’s native phospholipid composition. **2) Generation of chemically enriched OxPL reference pools.** In parallel, a subset of the biological sample is subjected to two oxidation reactions, Fenton (Fe^2+^/ H_2_O_2_) and H_2_O_2_-only. Each oxidation reaction is sampled at multiple time points to capture OxPLs across a range of oxidation depths and abundances.^37, 59^ Reference samples are profiled by LC-MS/MS on the same platform as the biological sample, providing empirical MS/MS spectra and retention-time anchors for OxPL features at concentrations sufficient for confident structural annotation. **3) Construction of a sample-specific *in silico* OxPL spectral library.** Non-oxidized phospholipid annotations from Step 1 are passed to LPPtiger 2,^43, 60^ which performs *in silico* oxidation to generate predicted OxPL structures and their theoretical MS/MS spectra. The library is therefore constrained to plausible OxPLs derived from the phospholipid species actually present in the sample, focusing the annotation search on the most biologically relevant species and minimizing false annotation.^33, 43^ **4) Reference-assisted OxPL annotation.** OxPLs in the biological sample are identified when they match (i) the retention time and *m/z* of an OxPL feature in the experimental reference pool, and (ii) the predicted MS/MS spectrum from the sample-specific *in silico* library. Critically, by including supporting MS/MS spectra from the reference pool when the species is below the MS/MS-quality threshold in the biological sample itself, RISOP recovers annotations for OxPLs that conventional untargeted lipidomics would miss.

### Oxidized Phospholipids Identification in Fenton and H_2_O_2_-only References

We used Fenton reaction and H_2_O_2_-only treatment to generate oxidized phospholipid (OxPL) references. While most previous studies have used reaction times of 12 to 24 hours,^37, 59^ we included multiple time points for both reaction types to capture a broader range of oxidation products in different intensities. We then performed untargeted lipidomic analysis by LC-MS/MS to obtain the global lipidome of oxidized and non-oxidized lipids in all reference samples (**Figure 2A, Table S3, and Table S4**). Our lipidomics platform demonstrates analytical precision and system stability, which ensures accurate retention time and *m/z* matching required for RISOP (**Figure S1**). We identified a combined number of 79 OxPLs from Fenton reaction and H_2_O_2_-only references by using the sample-specific OxPL spectral library generated in these references.

**Figure 2.**
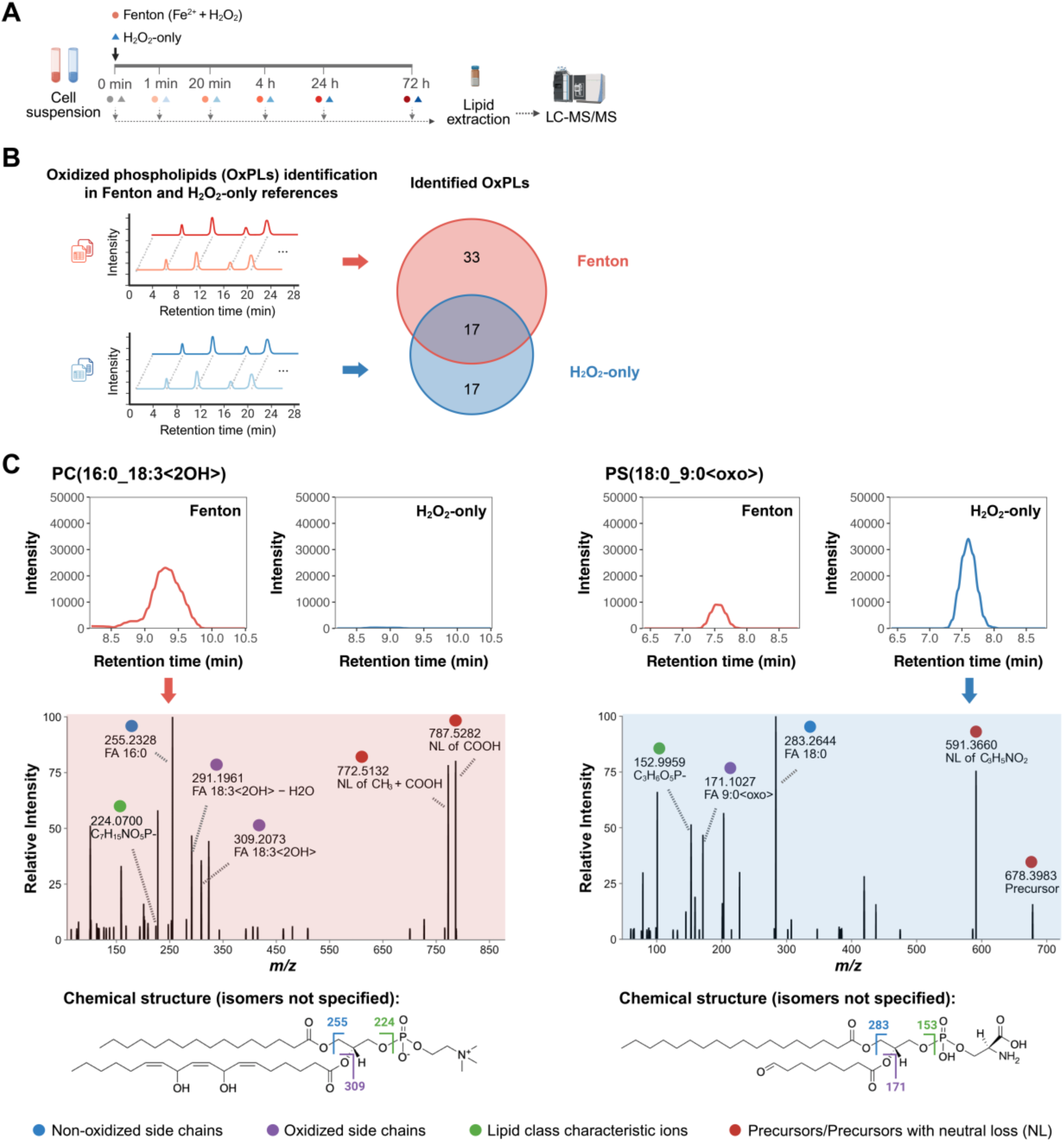
Identification of oxidized phospholipids in Fenton and H_2_O_2_-only references. (**A**) Preparation of OxPL references. KP4 cell suspensions were subjected to Fenton reaction and H_2_O_2_-only treatment for 5 different reaction time points. At the end of each reaction time point, samples were harvested for lipidomic analysis by LC-MS/MS. (**B**) OxPL identification in separate Fenton and H_2_O_2_-only references. Venn diagram showing the number of unique and commonly identified OxPLs from Fenton and H_2_O_2_-only references. (**C**) Representative Fenton-unique and H_2_O_2_-only-unique OxPLs. Top: Extracted ion chromatograms (EIC) of two representative OxPLs PC(16:0_18:3<2OH>) (left) and PS(18:0_9:0 <oxo>) (right) from Fenton references (red) and H_2_O_2_-only references (blue). Middle: Representative tandem mass spectra with diagnostic ions for identification. Each fragment ion is annotated with mass-to-charge ratio (*m/z*) and corresponding fragment identity or elemental composition. Fragments labeled in blue, purple, green, and red represent non-oxidized side chains, oxidized side chains, lipid class characteristic ions, and precursors or precursors with neutral loss (NL), respectively. Bottom: chemical structures of PC(16:0_18:3<2OH>) (left) and PS(18:0_9:0 <oxo>) (right) with isomers not specified. Annotations follow the identical color codes as in middle panel.

Importantly, because many OxPLs are of low abundance, we excluded species that were not reproducibly detected across replicates to minimize false annotation of background noise (coefficient of variation < 20%, see Methods), leaving 74 OxPLs for further analysis. In addition, oxidized lipids are generally labile and can undergo in-source fragmentation (ISF) – an artifactual fragmentation introduced during data acquisition that does not represent the true biological sample.^60, 61^ These undesired ISF products often exhibit identical or longer retention times compared to their non-oxidized counterparts, contributing another source of false annotation.^60–62^ To minimize misannotation of OxPLs from ISF products, identified OxPLs were visualized using retention time versus hydrogen-based Kendrick mass defect (H-KMD) plots.^60^ This approach allowed us to identify and exclude 7 potential ISF products (based on retention time, see Methods) from the 74 OxPLs in oxidized lipid references (**Figure S2**).

To compare OxPLs generated by each oxidation chemistry, Fenton and H_2_O_2_-only references were first analyzed independently. Of the 67 OxPLs identified across the two reference types, 33 were unique to Fenton, 17 were unique to H_2_O_2_-only, and only 17 were annotated in both (**Figure 2B**). Two distinct sources contribute to this limited overlap. First, the two chemistries generate partially non-overlapping product repertoires. For example, PC(16:0_18:3<2OH>) was identified only in Fenton references (**Figure 2C**), consistent with the more aggressive radical chemistry of the Fenton reaction. Second, OxPLs present at low abundance in one reaction type may fail to yield interpretable MS/MS spectra and remain unannotated even when the OxPL feature itself is detected. For instance, PS(18:0_9:0 <oxo>) was detected as an OxPL feature in both reference types but lacked MS/MS spectra in the Fenton-derived references, where its abundance was lower (**Figure 2C**). The latter source of missed annotations is analytical rather than chemical and motivates the integrated reference-assisted analysis of RISOP.

### Reference-assisted Combined Analysis on OxPL References Expands Identification Coverage and Dynamic Range of OxPLs

To test whether reference-assisted analysis could enhance OxPL identification beyond what either reference type yields alone, we analyzed all reference samples from Fenton and H_2_O_2_-only treatment together (**Figure 3A**). In this integrated analysis, an MS/MS spectrum acquired in one reaction type is permitted to support annotation of a feature at matching retention time and *m/z* in the other reaction type, provided the spectrum also matches a predicted spectrum from the sample-specific *in silico* OxPL library. This addresses a key limitation observed in the separate analyses (**Figure 2B**): many OxPL features are present in both reaction types but fall below the abundance threshold required to generate interpretable MS/MS spectra in one of them, and are therefore left unannotated when each reference is analyzed in isolation.

**Figure 3.**
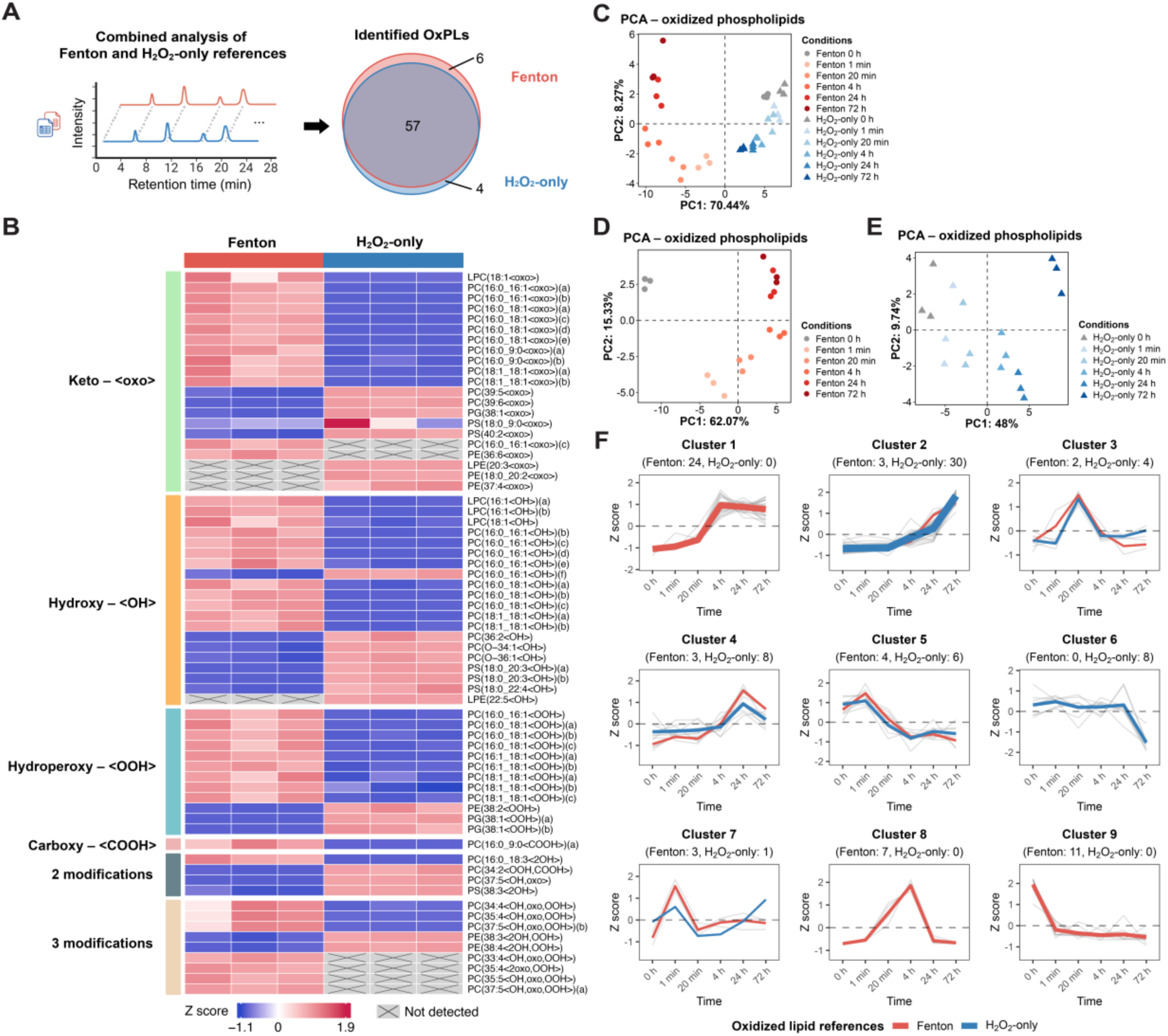
Reference-assisted combined analysis on OxPL references expands identification coverage and dynamic range of OxPLs. (**A**) OxPLs identified from combined analysis of Fenton and H_2_O_2_-only references. Venn diagram showing the number of unique and commonly identified OxPLs from Fenton and H_2_O_2_-only references. (**B**) Heatmap of 67 OxPLs identified in Fenton and/or H_2_O_2_ references. Each row represents one OxPL, and each column represents individual replicate of the Fenton or H_2_O_2_-only references. Columns are grouped by reference type, and rows are grouped by distinct oxidative modifications. Z scores representing the relative abundance of each OxPL are calculated from normalized OxPL intensity and plotted. Grey squares with crosses indicate OxPLs that were not detected. (**C**) Principal component analysis (PCA) on log2-transformed normalized intensity of 57 common OxPLs identified in Fenton (dots in shades of red and grey) and H_2_O_2_-only (triangles in shades of blue and grey) references from untargeted lipidomics. Each data point represents one sample at a specific time point from either Fenton or H_2_O_2_-only references. (**D-E**) PCA on log2-transformed normalized intensity of 63 OxPLs identified in Fenton (**D**) and 61 OxPLs identified in H_2_O_2_-only (**E**) references from untargeted lipidomics. Each data point represents one sample at a specific time point from either Fenton references or H_2_O_2_-only references. (**F**) K-means clustering on 57 common OxPLs identified in both Fenton and H_2_O_2_-only references. K-means clustering was performed on normalized intensity of OxPLs in reference samples from different time points of Fenton or H_2_O_2_-only treatment. Number of OxPLs from Fenton or H_2_O_2_-only references in each cluster is indicated. Z scores of individual OxPLs (grey lines) are shown together with red and blue lines representing the mean Z score from Fenton and H_2_O_2_-only references, respectively. Line thickness is proportional to the number of OxPLs. Dashed line indicates a Z score of zero.

Consistent with this rationale, integrated analysis yielded 57 OxPLs annotated in both reference types, representing a marked increase from the 17 overlapping species recovered by separate analysis (**Figure 2B**) without changing the union of 67 identified OxPLs (**Figure 3A**). The expanded overlap therefore reflects analytical recovery of true co-occurrence rather than a change in the underlying OxPL repertoire, demonstrating that MS/MS sharing between reaction types within RISOP rescues annotation of low-abundance OxPLs that would otherwise be missed. Beyond identification count, the two reaction types differed in the types of oxidative modification they enriched. Among the 67 identified OxPLs, species carrying hydroperoxy modifications (<OOH>), products of the early oxidation cascade,^5, 6^ were relatively more abundant in Fenton references. Fenton references also contained a considerably larger fraction of OxPLs bearing three oxidative modifications per molecule (**Figure 3B**), consistent with the high reactivity of hydroxyl radicals generated in this system and indicating that Fenton chemistry promotes more extensive per-molecule oxidation. Together, these results show that RISOP’s reference-assisted approach substantially expands OxPL annotation while preserving informative differences in modification profile between the two oxidation chemistries.

Enhanced OxPL identification enables global OxPL analyses that are otherwise restricted to a limited number of OxPLs. Principal Component Analysis (PCA) on OxPLs revealed that Fenton reference samples with varying reaction time formed distinct clusters that were clearly separated from those of the H_2_O_2_-only references (**Figure 3C**). In addition, oxidized reference samples were well separated by reaction time along principal component 1 and 2, suggesting progressive temporal changes in OxPL composition under both Fenton and H_2_O_2_-only oxidation conditions when all samples were plotted together (**Figure 3C**), or separately by reaction type (**Figure 3D, 3E**). PCA on over 710 non-oxidized lipids largely recapitulated clustering by oxidation reaction and reaction time (**Figure S3A-S3C**). Thus, global OxPL analysis reveals time-dependent, progressive changes in OxPL profiles under both oxidation conditions.

To further examine changes of OxPL at individual lipid level, we performed K-means clustering on identified OxPLs to uncover groups of OxPLs that exhibit similar temporal trajectory.^63, 64^ Though a few clusters capture lipids with similar time-dependent changes in intensity from both types of references, the majority of identified clusters reveal distinct patterns that are predominantly found in either Fenton or H_2_O_2_-only references (**Figure 3F**). Specifically, clusters 1, 8, and 9 were mostly associated with the Fenton reaction, whereas clusters 2 and 6 were enriched under H_2_O_2_-only oxidation, indicating divergent OxPL formation dynamics between these two types of oxidation reactions. Thus, these results indicated that oxidation reactions of Fenton and H_2_O_2_-only treatments produce OxPLs with different temporal patterns (**Figure 3F**). Together, a reference pool created with 2 oxidation reactions and 5 time points expands the coverage and dynamic range of OxPLs that will aid in enhanced identification of OxPLs in biological samples.

### Applying RISOP in Studying Ferroptosis-associated Lipid Peroxidation

To demonstrate the application of RISOP to studying lipid peroxidation, we applied this method to investigate ferroptosis, an iron-dependent cell death characterized by hallmark accumulation of lipid peroxidation.^65, 66^ We used KP4 cells, a human pancreatic ductal cell carcinoma cell line with known susceptibility to ferroptosis as our model.^67^ We verified that the viability of KP4 cells is sensitive to ferroptosis inducer ML210 in a dose-dependent manner, and can be rescued by co-treatment with ferrostatin-1 (Fer-1), a selective inhibitor of ferroptosis (**Figure S4A**).^68^

We harvested KP4 cells treated with ML210 at 6 different time points for lipidomic analysis (**Figure 4A, Table S5**). The dose and time points were chosen at which cells had not yet undergone ferroptosis (**Figure S4B**), but may have already accumulated different levels of lipid peroxidation. We first compared the number of OxPLs identified in ML210-treated samples alone in the absence of any references, or from combined analyses using one or both types of OxPL references through RISOP (**Figure 4B**). As expected, in the absence of Fenton or H_2_O_2_-only references, only 9 OxPLs were identified in ML210-treated cells. Incorporation of reference samples obtained from either Fenton or H_2_O_2_-only treatment substantially increased identification of OxPLs. Notably, the combined use of all references more than doubled the number of OxPLs from 9 to 22, demonstrating the power of RISOP in enhancing the identification of oxidized lipid species in biological samples (**Figure 4B**). The molar concentrations of OxPLs quantified in our samples were in good agreement with previously reported endogenous levels of oxidized phospholipids in mammalian cells and tissues.

**Figure 4.**
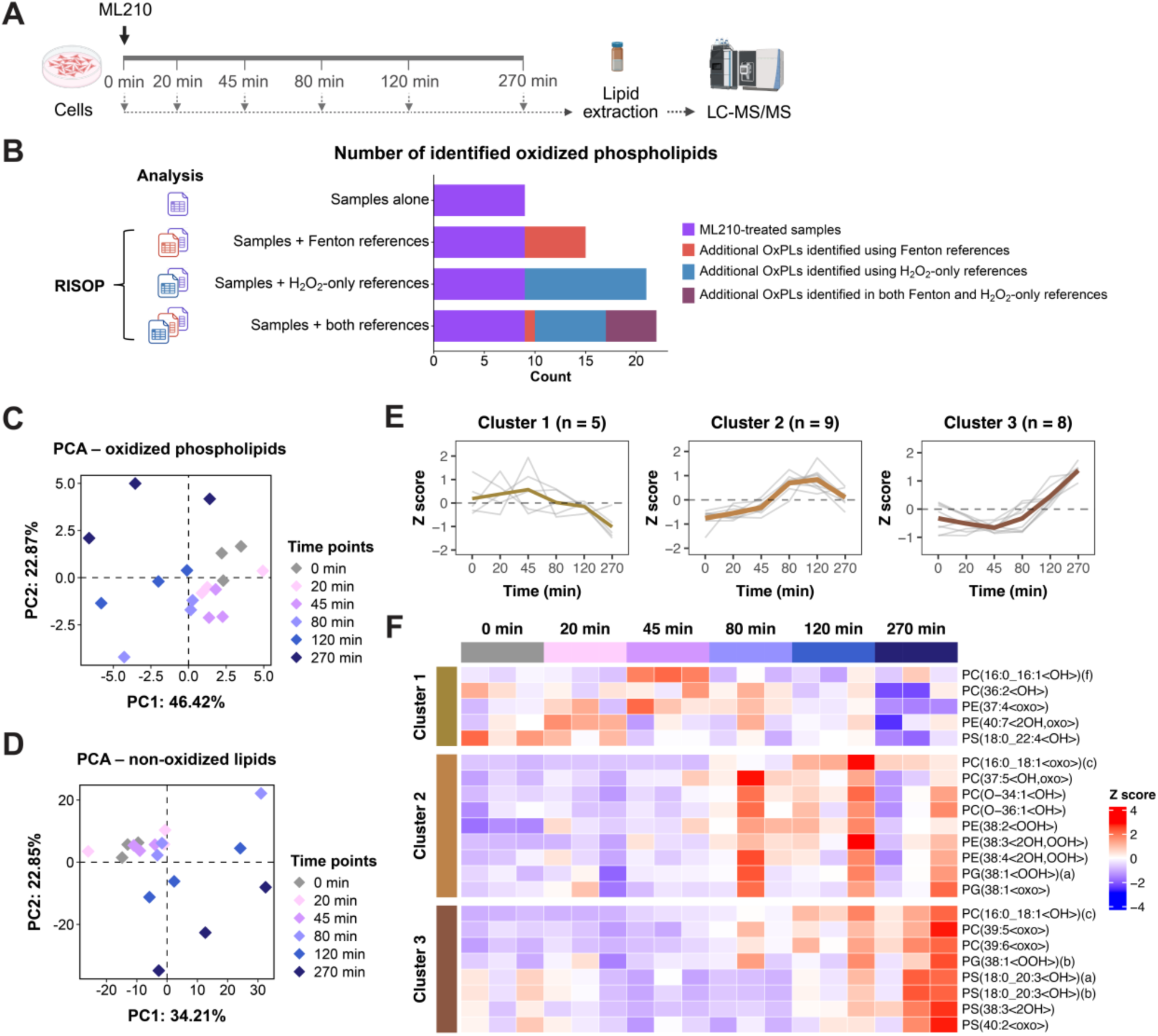
Applying RISOP in studying ferroptosis-associated lipid peroxidation. (**A**) Preparation of ML210-treated cells. KP4 cells were incubated with 10 μM ML210 for six treatment time points. At the end of each incubation time point, samples were harvested for lipidomic analysis by LC-MS/MS. (**B**) Number of identified OxPLs in ML210-treated samples with or without RISOP. Stacked bar plot showing numbers of OxPLs identified in ML210-treated samples alone without RISOP (purple), and additional OxPLs identified using Fenton references (red), H_2_O_2_-only references (blue), or both Fenton and H_2_O_2_-only references combined (dark magenta) through RISOP. (**C**) Principal component analysis (PCA) on log2-transformed molar concentrations of 22 OxPLs identified in ML210-treated samples. Each data point represents one sample harvested from a given time point. (**D**) PCA on log2-transformed molar concentrations of 723 non-oxidized lipids identified in ML210-treated cells. Each data point represents one sample harvested from a given time point. (**E**) K-means clustering on 22 OxPLs identified in ML210-treated samples. K-means clustering was performed on molar concentrations of OxPLs in ML210-treated samples treated for different durations. In each cluster, number of OxPLs is indicated. Z scores of individual OxPLs (grey lines) are shown together with brown lines in different shades representing the mean Z score of each cluster. Line thickness is proportional to the number of OxPLs. Dashed line indicates a Z score of zero. (**F**) Heatmap of 22 OxPLs identified in ML210-treated samples. Each row represents one OxPL, and each column represents one biological replicate at the indicated ML210 treatment time point. Columns are grouped by treatment time and ordered from 0 to 270 min. Rows are grouped according to the K-means clusters defined in Figure 4E. Row-wise transformed Z scores of molar concentrations of OxPLs from each identified cluster are presented. Only OxPL isomers (indicated with letter suffixes in parentheses) that are identified in ML210-treated samples were included.

Specifically, our total OxPC (0.19-0.37 nmol/mg protein) and total OxPE (0.09-0.15 nmol/mg protein) are comparable to the total OxPC and OxPE quantified in mouse peritoneal macrophages by LC-MS/MS (after protein-per-cell conversion based on typical mammalian cells),^33^ which fall within the same sub-nmol/mg protein order of magnitude. Expressed as a fraction of the parent non-oxidized phospholipid pool, our total OxPC/total PC (0.09-0.13 mol%) and total OxPE/total PE (0.27-0.34 mol%) ratios fall within the <1–2 mol% range typically associated with endogenous, non-pathological membrane oxidation.^50, 69^ Collectively, these comparisons indicate that OxPLs detected in our study are consistent with previously reported endogenous oxidized phospholipid content in mammalian cells. To assess changes in the global lipidome, we performed PCA on all OxPLs identified in ML210-treated cells (**Figure 4C**). Our results revealed some separation in cells with longer ML210 treatment, indicating that ML210 induces changes in OxPL composition. Notably, analysis of non-oxidized lipids exhibited a similar clustering with increasing treatment time, suggesting that an elevated lipid peroxidation burden is accompanied by remodeling of the native, non-oxidized lipid environment as well (**Figure 4D**).

We next performed K-means clustering to further characterize the temporal changes of OxPL. We observed that OxPLs could be classified into three distinct clusters based on their temporal trajectories, each exhibiting a distinct pattern over time. Specifically, Cluster 1 remained relatively stable at early time points (0 – 45 min) and then decreased at later time points (≥80 min). Cluster 2 increased over time, peaked at 120 min, and then slightly declined. Cluster 3 showed an initial decrease followed by a marked increase, reaching the highest level at 270 min (**Figure 4E, 4F**). These results suggest that lipid peroxidation is a dynamic process, wherein elevated cellular oxidative burden leads to distinct responses in subgroups of OxPLs. Together, these results demonstrate that reference-assisted analysis using RISOP with oxidized lipid references markedly improves identification of OxPLs in biological samples undergoing increased oxidative stress.

### Lipid Peroxidation Assessment by BODIPY C11 and OxPL Profiling

Currently, one of the most widely used methods for assessing lipid peroxidation in live cells is the BODIPY C11 probe, which detects lipid radicals, a peroxidation intermediate, through oxidation of its polyunsaturated C11 moiety, leading to a shift in fluorescence emission from the reduced state (∼590 nm) to the oxidized state (∼510 nm).^70^ However, it remains unclear whether BODIPY C11 accurately reflects the overall burden of lipid peroxidation in living cells. To address this question, we used the same treatment paradigm that we established earlier (**Figure 4A**) and performed BODIPY C11 staining by flow cytometry (**Figure 5A and Figure S5A**). The level of cellular peroxidation assessed by BODIPY C11 increased and plateaued in cells treated with ML210 for 120 min, before decreasing to a similar level as a shorter treatment time (**Figures 5B, 5C and Figure S5B**). To compare changes in BODIPY C11 peroxidation with the level of total OxPL abundance by mass spectrometry, we performed correlation analysis between the percentage of peroxidation-positive cells and the total abundance of OxPLs along the course of treatment times. Interestingly, the proportion of peroxidation-positive cells did not correlate with total OxPL abundance in cells (**Figure S6**), suggesting that the BODIPY C11 signal may not reflect changes in overall OxPL abundance. However, when the percentage of peroxidation-positive cells was compared to each distinct temporal OxPL cluster that we identified (**Figure 4E**), a significant positive correlation was observed specifically with Cluster 2 (**Figure 5D**, *P* = 0.004, Pearson correlation followed by Benjamini–Hochberg correction).

**Figure 5.**
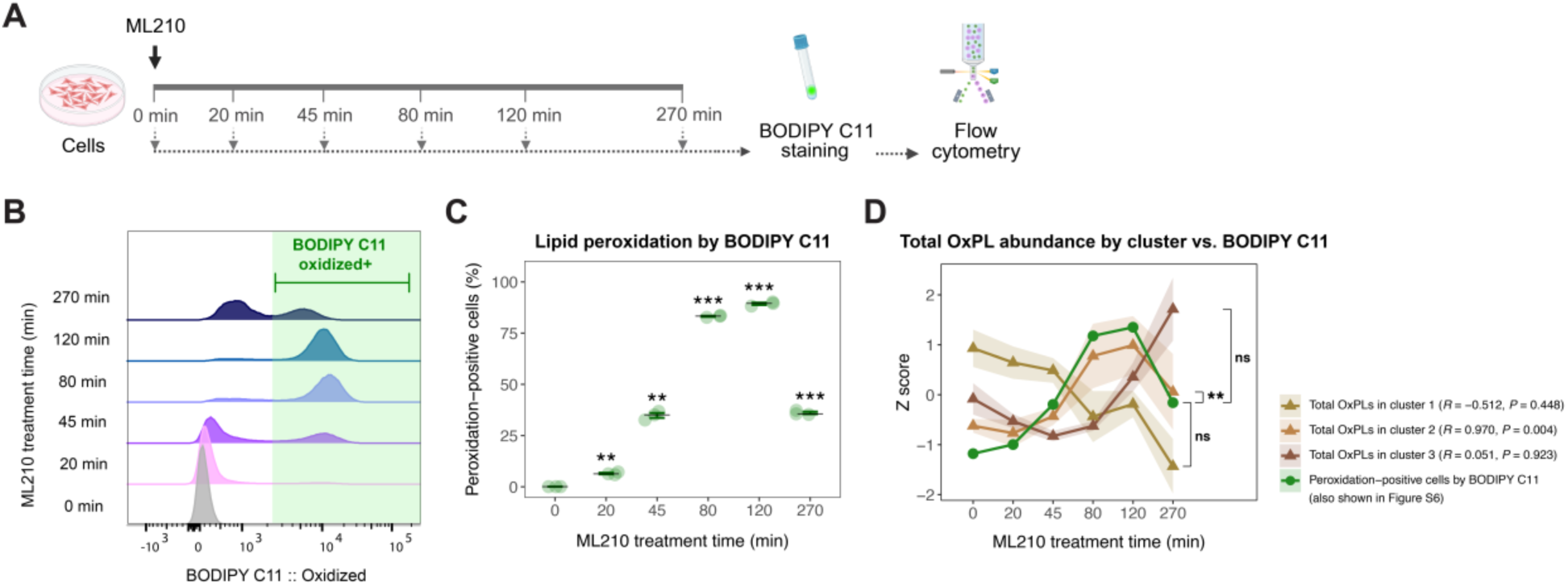
Lipid peroxidation assessment by BODIPY C11 and OxPL profiling. (**A**) Preparation of ML210-treated cells. KP4 cells were incubated with 10 μM ML210 for six treatment time points. At the end of each incubation time point, samples were harvested for BODIPY C11 staining followed by flow cytometry analysis. (**B-C**) Lipid peroxidation assessed by BODIPY C11 in samples treated with ML210. **B**, Histograms of BODIPY C11 staining intensity in FITC channel (515/25 nm, representing oxidized form) by flow cytometry are shown. Area shaded in green indicates peroxidation-positive cells. **C**, Dot-and-whisker plot showing quantification of peroxidation-positive cells. Each replicate (individual dots) is shown with error bars representing mean ± SEM (n = 3). Asterisk indicates significance compared to the 0 min control (\*\*\**P* < 0.001, \*\**P* < 0.01; Welch’s *t*-test with Benjamini–Hochberg adjustment). (**D**) Total OxPL abundance by cluster vs. BODIPY C11 lipid peroxidation assay. Total OxPL intensity of each defined cluster (from Figure 4E) was calculated and transformed into Z score. Z score of each cluster is plotted (lines in shades of brown) together with Z score of the percentage of peroxidation-positive cells by BODIPY C11 staining (Figure 5C) (green line, also shown in Figure S6). Shaded area indicates SEM (n = 3) on all lines. Pearson correlation analysis was performed to assess the relationship between time-dependent changes in peroxidation-positive cells and total OxPL abundance in each cluster. Correlation coefficient (*R*) and BH-adjusted *P*-value of each correlation are shown (\*\**P* < 0.01; ns, non-significant). Abstract Graphic

Cluster 2 contains a larger number of OxPLs with hydroperoxy modifications (<OOH>) than the other 2 clusters (**Figure 4F**). Given that lipids with hydroperoxy modifications (<OOH>) are the direct product of the same radical chemistry that BODIPY C11 reacts to,^70^ it is therefore plausible that the abundance of OxPLs in Cluster 2 correlates with the readout of BODIPY C11. These results suggest that BODIPY C11 may only capture the oxidative burden of a subset of OxPLs in live cells. The lack of a global correlation between BODIPY C11 and total OxPL abundances likely arises from their fundamentally different readouts. Specifically, BODIPY C11 reports ongoing radical-mediated lipid peroxidation,^71^ whereas lipidomics-based OxPL analysis captures a broad spectrum of oxidized lipid products, including both early- and late-stage oxidation products. These results highlight the need for comprehensive identification of OxPLs in understanding their contribution to key cellular processes.

## CONCLUSIONS

In this study, we developed RISOP, a reference-assisted approach for enhanced identification of oxidized phospholipids (OxPLs) in an untargeted lipidomics workflow by LC-MS/MS. The core advance of RISOP is to generate chemically enriched OxPL reference pools from the same biological matrix using Fenton and H_2_O_2_-only oxidation, providing high-quality MS/MS spectra and retention-time anchors for OxPLs that are otherwise too scarce to be annotated directly. By sharing MS/MS evidence across the two reaction types and matching against a sample-specific *in silico* spectral library, RISOP recovers identifications for low-abundance OxPLs while constraining the search to biologically plausible species. This strategy more than doubled the number of OxPLs identified in ferroptosis-induced KP4 cells, without requiring modifications to instrumentation or data acquisition. Beyond method development, we found that the widely used BODIPY C11 lipid peroxidation probe captures oxidative burden in only a subset of OxPLs in live cells – underscoring the need for comprehensive, untargeted characterization of the OxPL landscape in biological processes. RISOP is generalizable and can be applied to studies of different cell and tissue types beyond cultured cell systems. Our method can be expanded to investigate other classes of oxidized complex lipids, including oxidized glycerol lipids, offering a broadly applicable platform for lipidomics studies. The improved identification capability established here lays the groundwork for future mechanistic investigations into the functional roles of oxidized complex lipids in physiology and disease.

## AUTHOR INFORMATION

### Author Contributions

Z.C. designed the project under the guidance of X.Z.. Z.C. performed all experiments and data analyses. A.E. conducted independent code checking and accuracy checks. A.E. and N.I. provided feedback on graphic design and the written manuscript. Z.C. wrote the manuscript with X.Z..

## Acknowledgements

This work was supported by a translational geroscience pilot award from the Yale Claude D. Pepper Older Americans Independence Center (X.Z.) and McCance Fellowship (X.Z.). We thank Dr. Mandar Muzumdar at Yale School of Medicine for generously providing the KP4 cell line used in this study. We thank Yale Flow Cytometry for their assistance with flow cytometry analysis. Schematic diagrams were created with BioRender.com.

## Code availability

The R scripts generated for this study can be accessed at: https://github.com/ZhaoLab-Yale/RISOP_R_code

## Supplementary Figures

**Figure S1.**
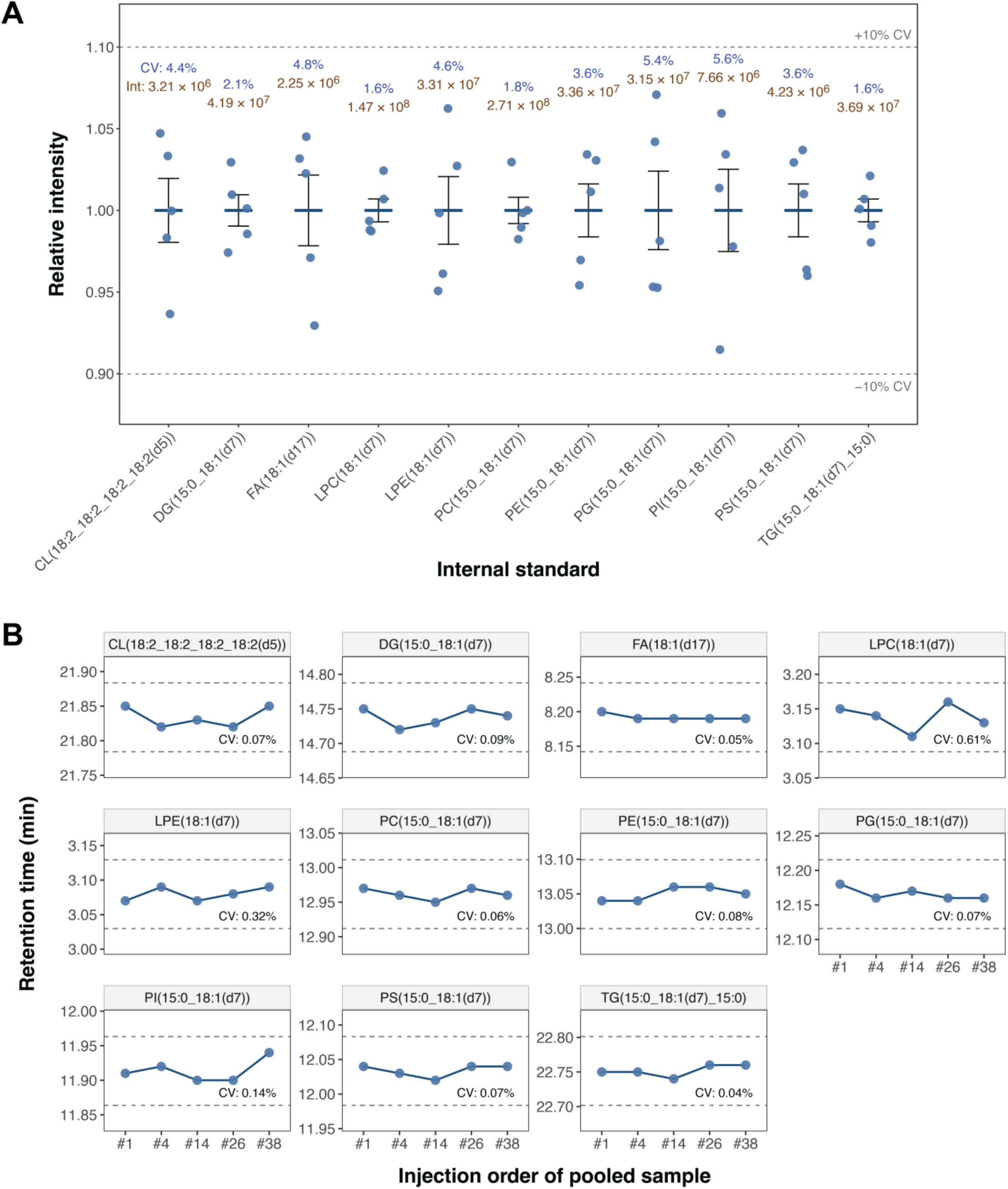
System stability and reproducibility. (**A**) Intensities of deuterated internal standards in pooled samples throughout this study. Dot-and-whisker plot showing the normalized intensity of each deuterated internal standard. Each injection (individual dots) is shown with error bars representing mean ± SEM (n = 3). Absolute mean intensity (Int; brown) and coefficient of variation (CV; blue) are included above each internal standard. Dashed lines indicate ±10% CV from the mean normalized intensity. (**B**) Retention time monitoring throughout this study. Retention time (Y-axis) of each deuterated internal standard is plotted against sequential injection order of the pooled sample (X-axis). CV of retention time is shown in each panel. Dashed lines indicate ±0.05 min from the mean retention time of each internal standard.

**Figure S2.**
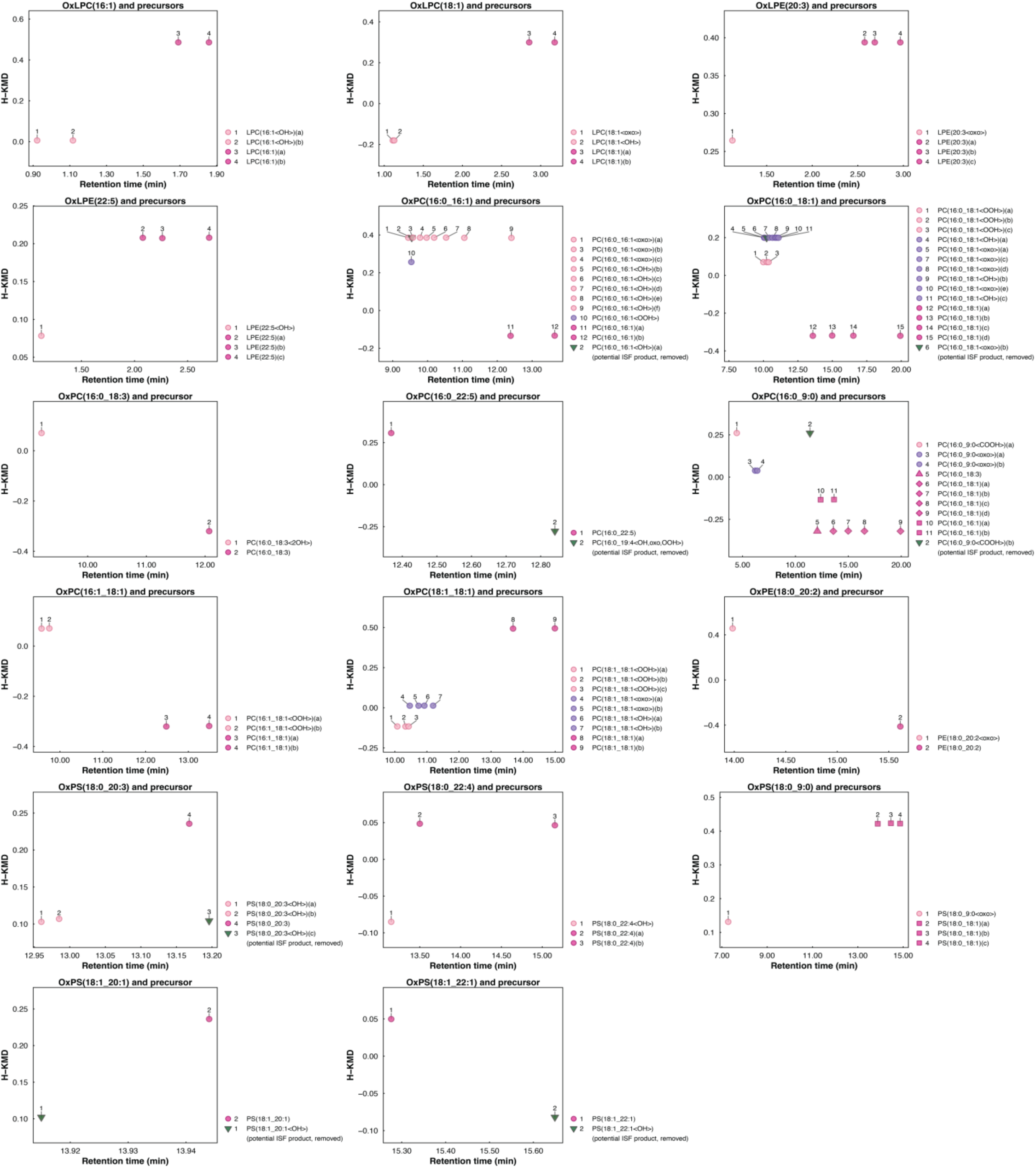
Hydrogen-based Kendrick mass defect (H-KMD) plots of identified oxidized phospholipids and their corresponding non-oxidized precursor(s). The X-axis represents the chromatographic retention time, and the Y-axis represents the H-KMD, calculated by scaling the measured *m/z* to the nominal mass of hydrogen. Each data point represents an individual lipid species. Non-oxidized precursor lipids are shown in magenta, and oxidized lipids are shown in pink or purple. Shapes indicate different OxPL and precursor pairings. Circles are used when OxPLs and their corresponding precursors share identical fatty acyl compositions, whereas distinct precursors with fatty acyl compositions different from those of product OxPLs are represented by squares, diamonds, or triangles. Potential in-source fragmentation (ISF) products (removed from data analysis) are shown as inverted triangles in green.

**Figure S3.**
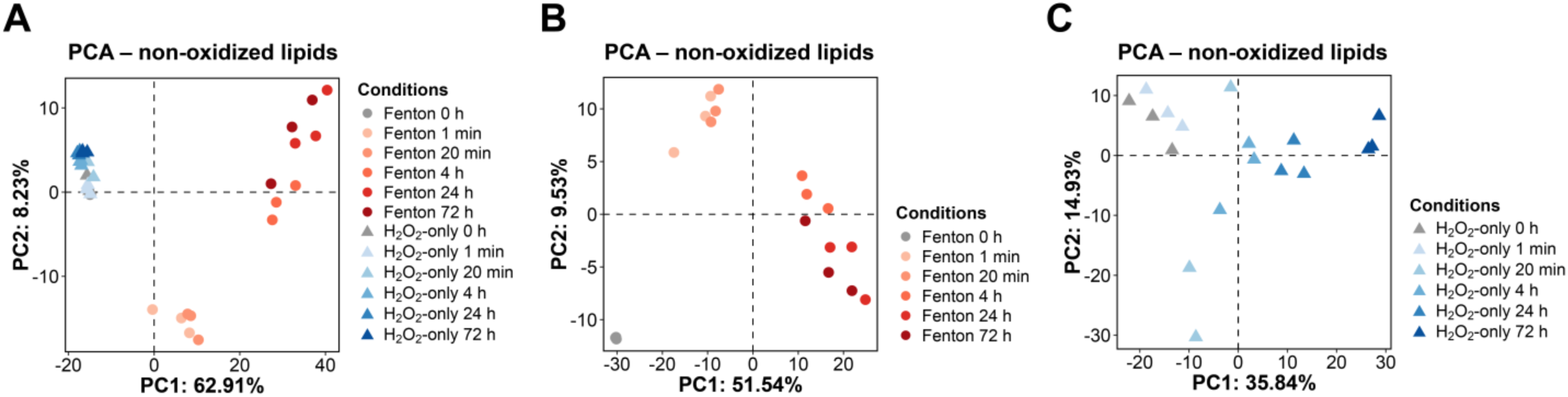
Non-oxidized lipidomic profiles of Fenton and H_2_O_2_-only references. (**A**) Principal component analysis (PCA) on log2-transformed normalized intensities of 715 common non-oxidized lipids identified in Fenton (dots in shades of red and grey) and H_2_O_2_-only (triangles in shades of blue and grey) references. Each data point represents one sample at a specific time point from either Fenton or H_2_O_2_-only references. (**B-C**) PCA on log2-transformed normalized intensities of 715 non-oxidized lipids identified in Fenton (B) and 739 non-oxidized lipids identified in H_2_O_2_-only (C) references. Each data point represents one sample at a specific time point from either Fenton or H_2_O_2_-only references.

**Figure S4.**
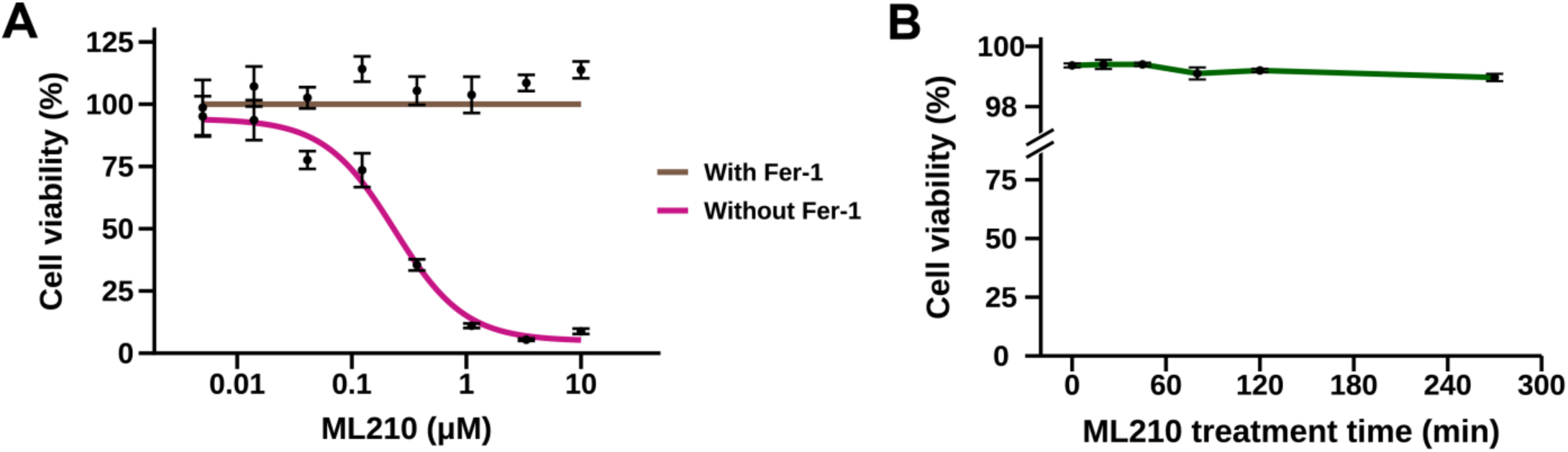
Cell viability with ML210 treatment. (**A**) Viability of KP4 cells treated with different concentrations of ML210 for 24 hours in the absence (magenta) or presence (brown) of ferrostatin-1 (Fer-1). Mean viability (dot) is presented together with error bars (± SEM) (n = 6). (**B**) Viability of KP4 cells treated with 10 μM of ML210 at 6 different time points. Mean viability (dot) is presented together with error bars (± SEM) (n = 3).

**Figure S5.**
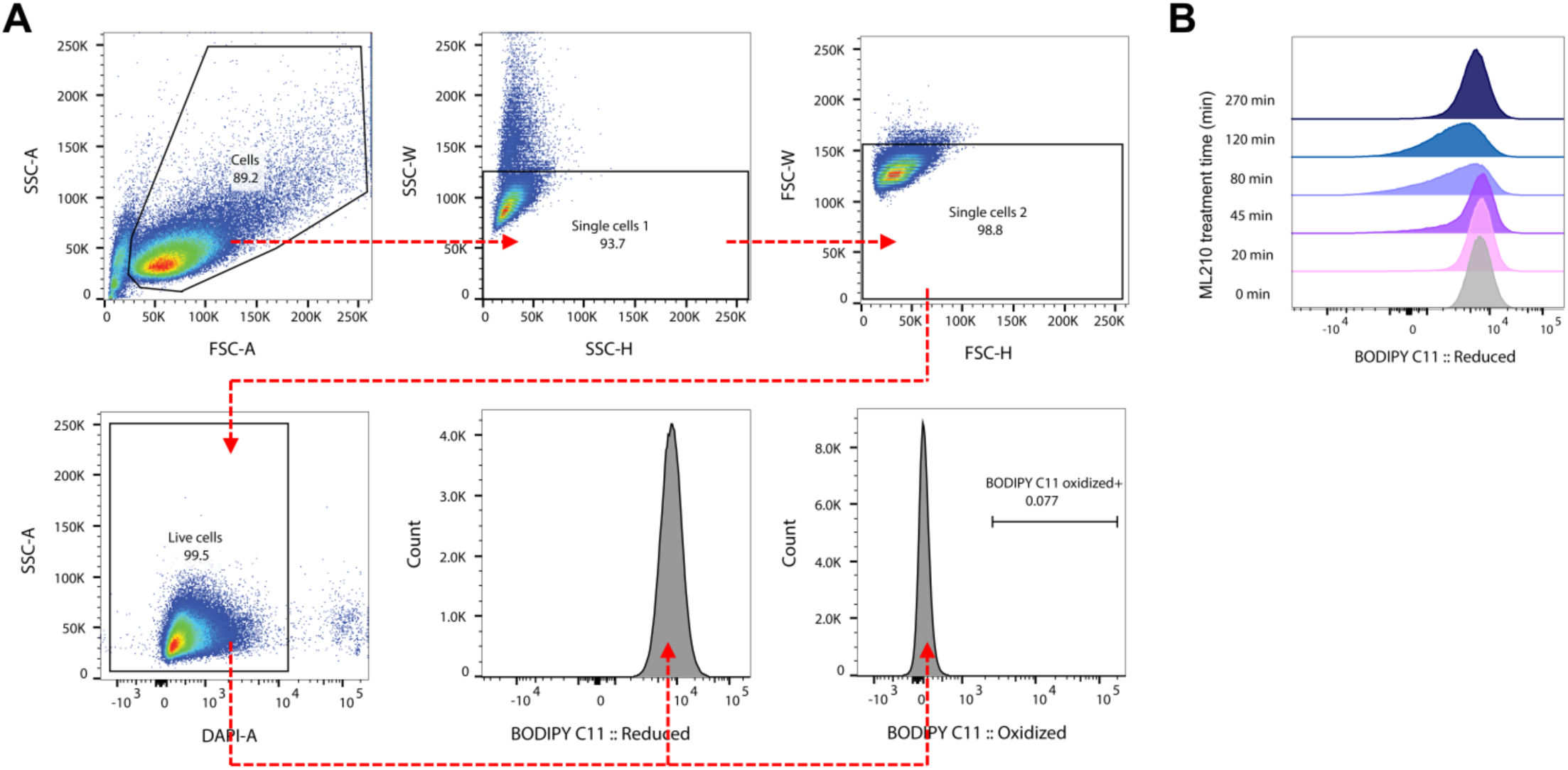
BODIPY C11 lipid peroxidation assay by flow cytometry. (**A**) FACS gating strategy used to quantify peroxidation-positive cells by BODIPY C11 staining. Cells without ML210 treatment were used as negative gating controls to determine the gate for BODIPY C11 peroxidation-positive cells. (**B**) Histograms of BODIPY C11 staining intensity in PE-Texas Red channel (610/20 nm, representing non-oxidized form) by flow cytometry are shown for ML210-treated samples.

**Figure S6.**
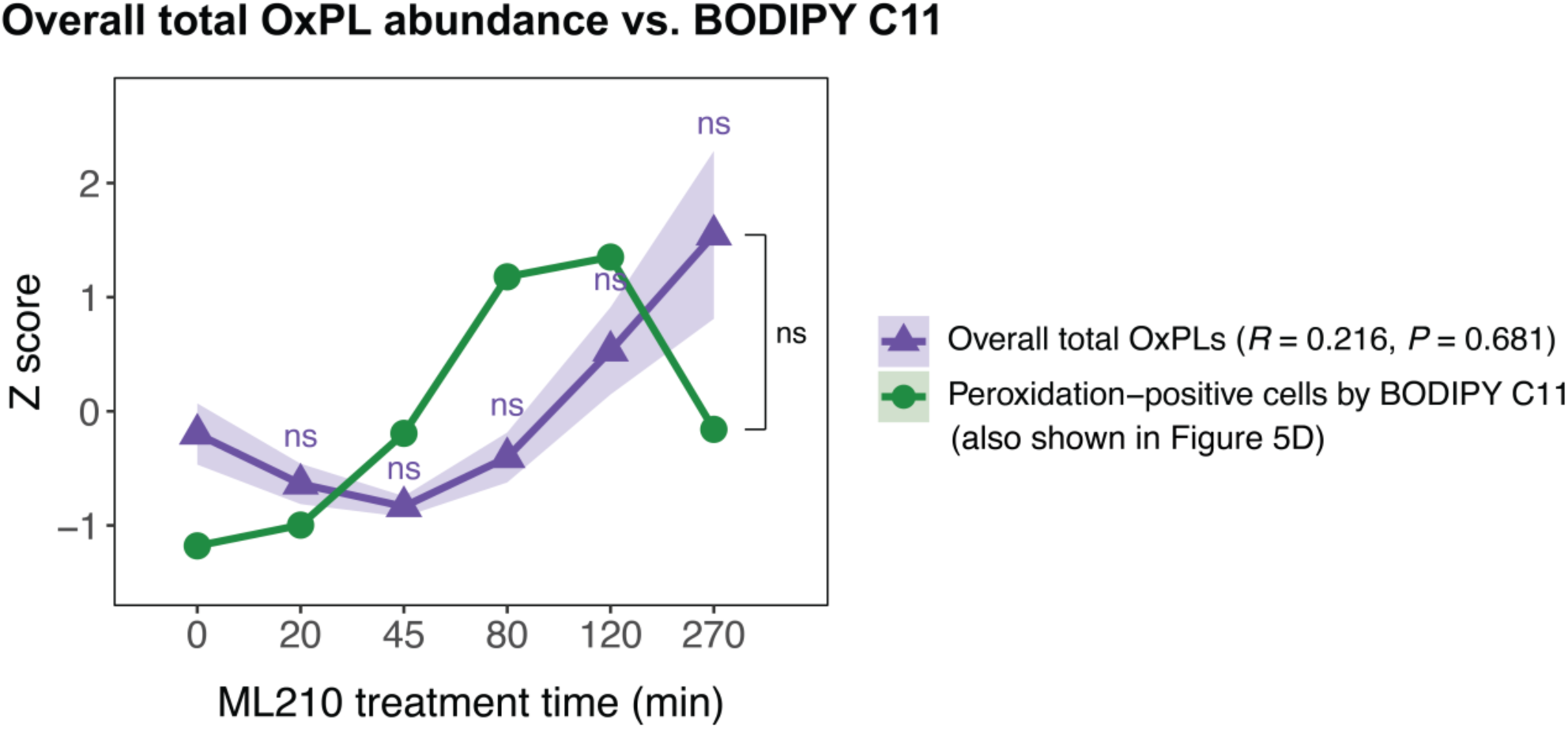
Overall total OxPL abundance vs. BODIPY C11 lipid peroxidation assay. Z scores of the percentage of peroxidation-positive cells by BODIPY C11 staining (green, also shown in Figure 5D) are plotted together with Z scores of total OxPL intensity (purple) in samples treated with ML210. Statistical significance at each time point was tested against 0 min control using Welch’s *t*-test with Benjamini–Hochberg adjustment (ns = non-significant). Shaded areas indicate SEM (n = 3). Pearson correlation analysis was performed to assess the relationship between time-dependent changes in peroxidation-positive cells and total OxPL abundance. Correlation coefficient (*R*) and *P* value are shown.

**Table S1.**
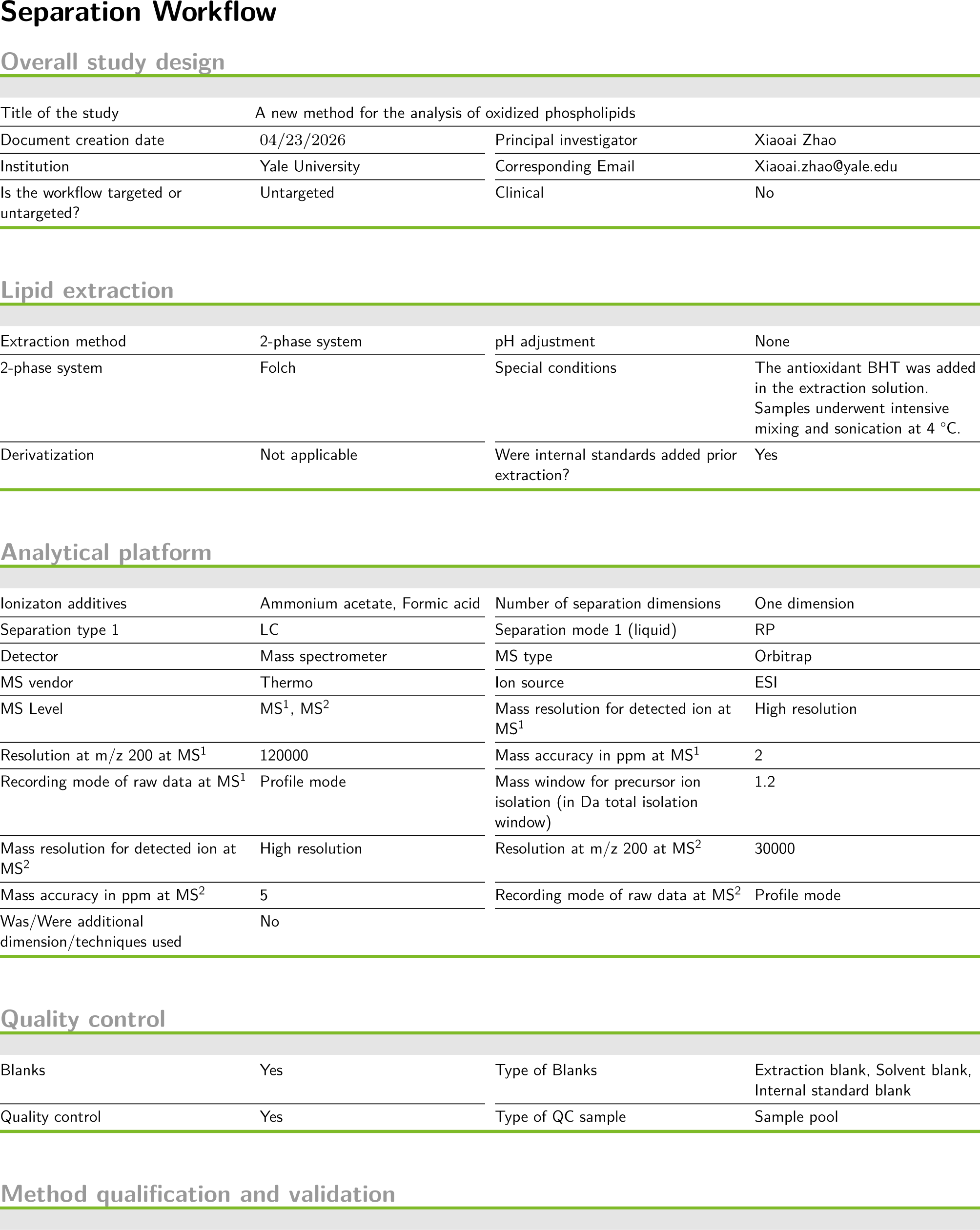
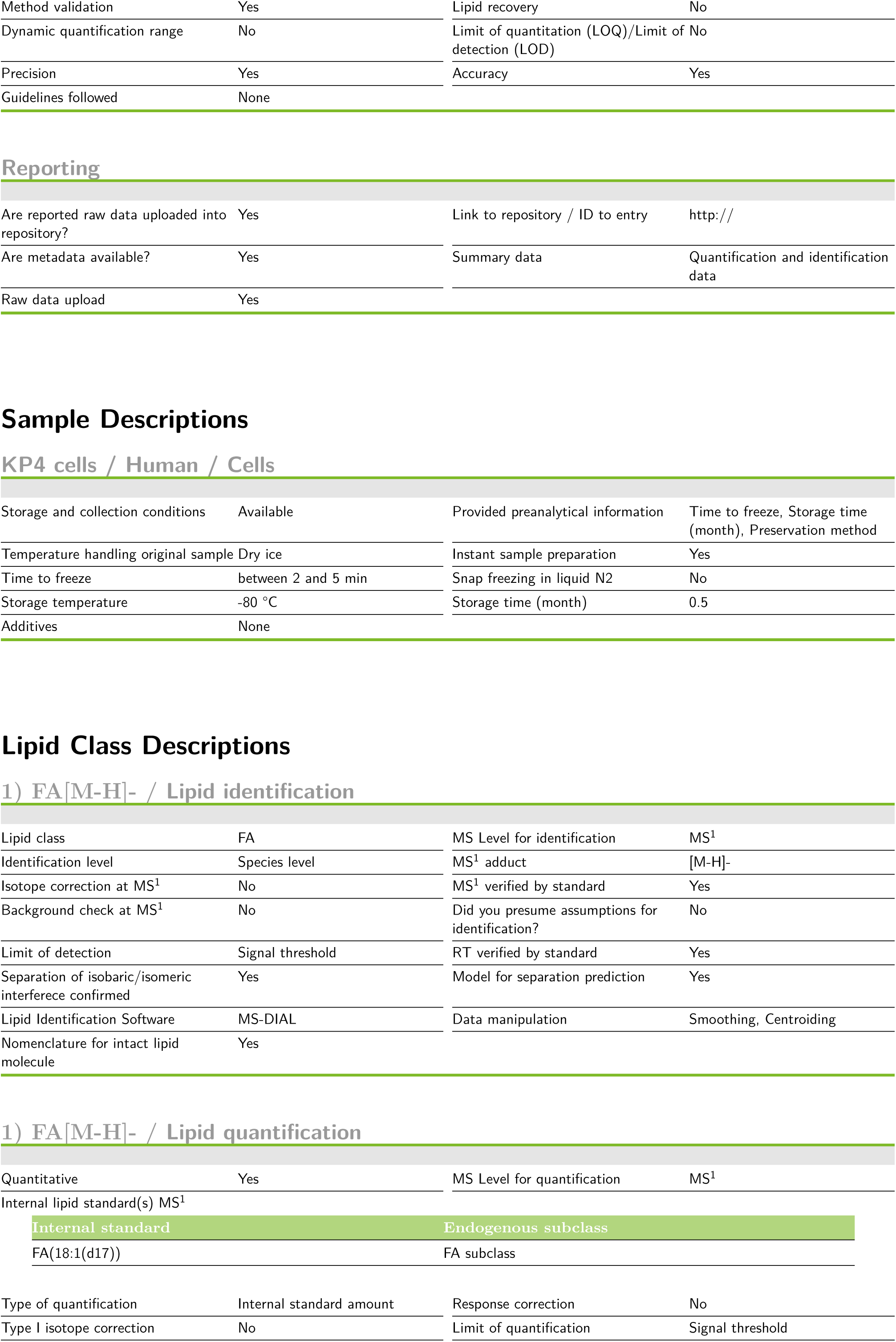
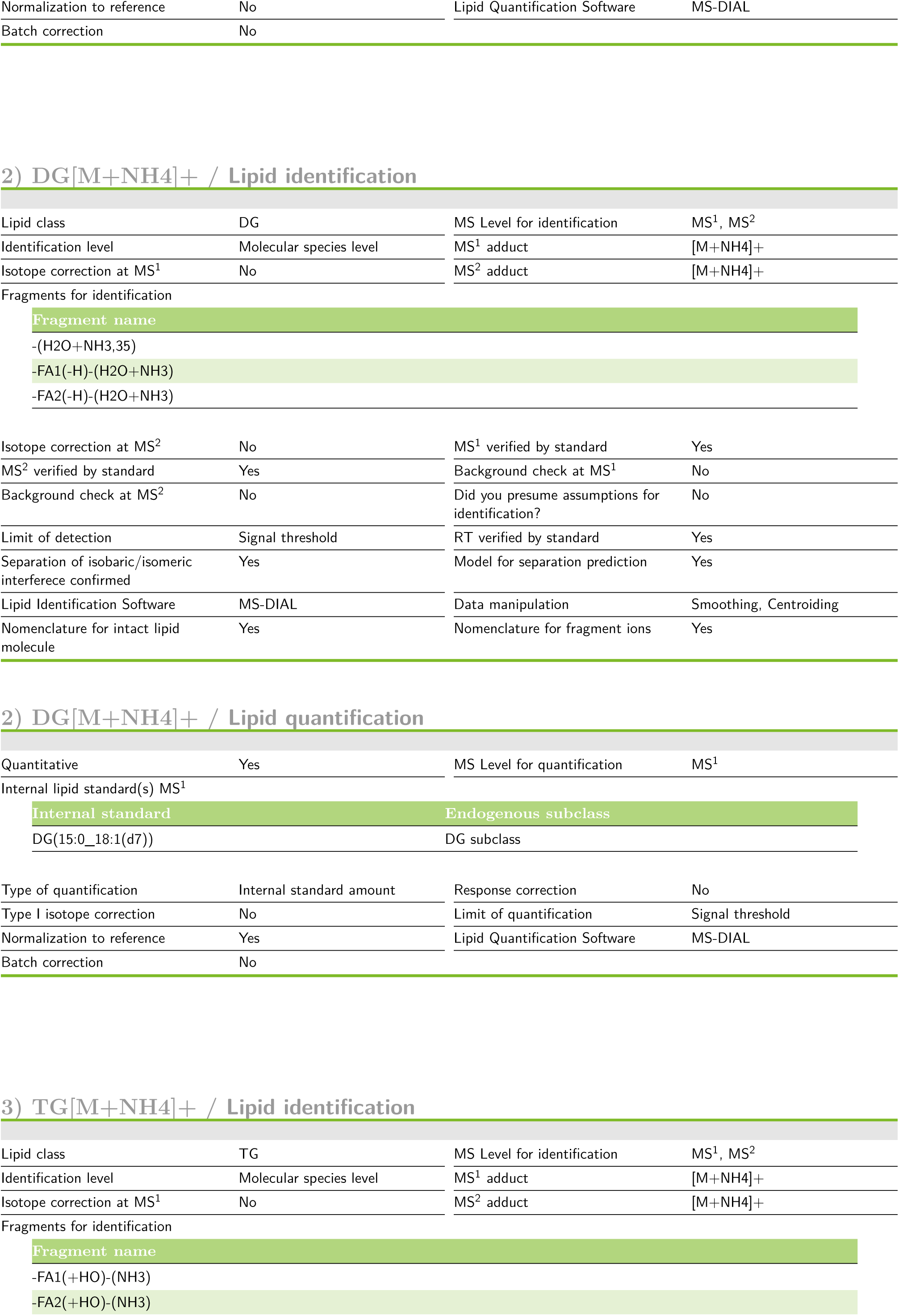
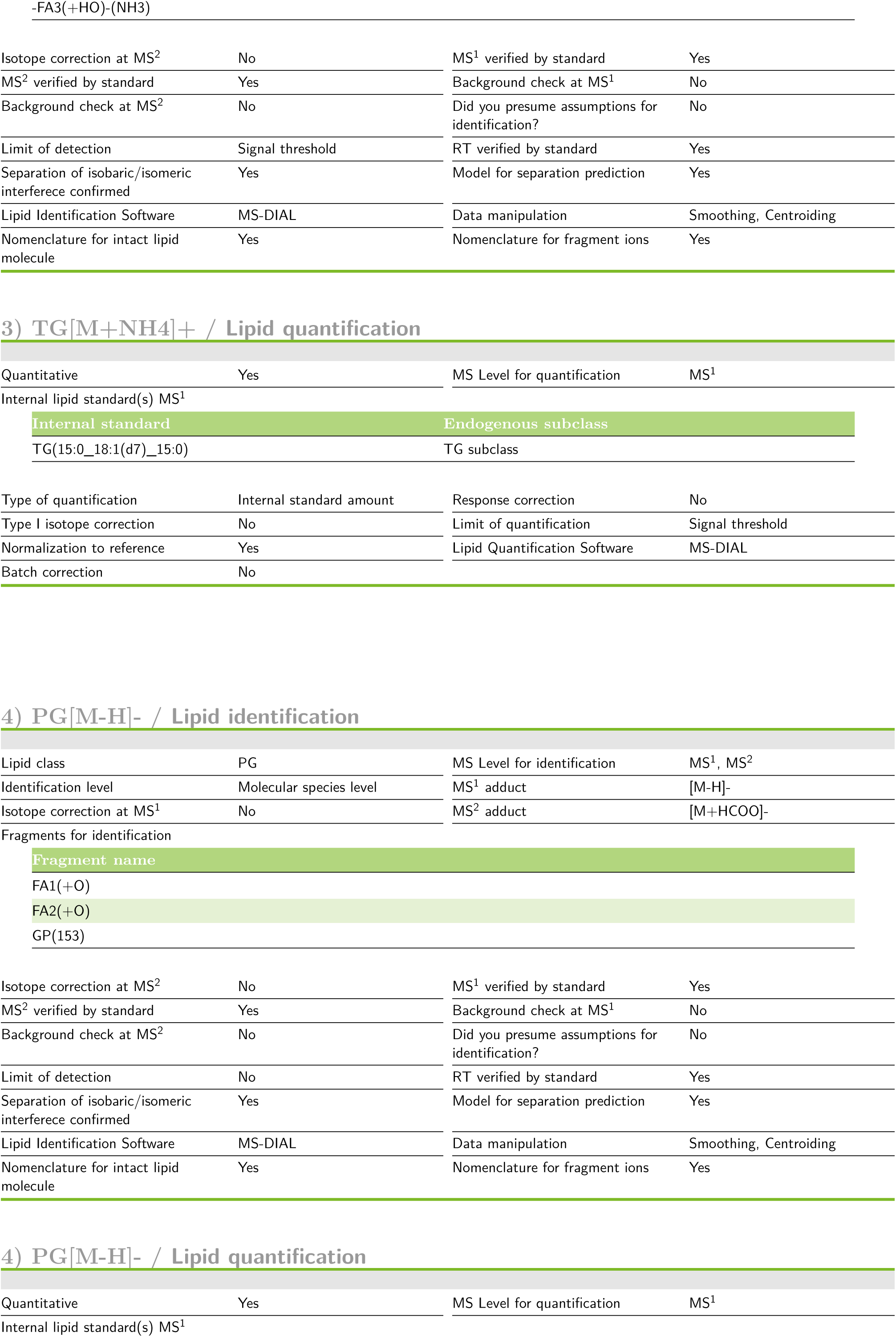
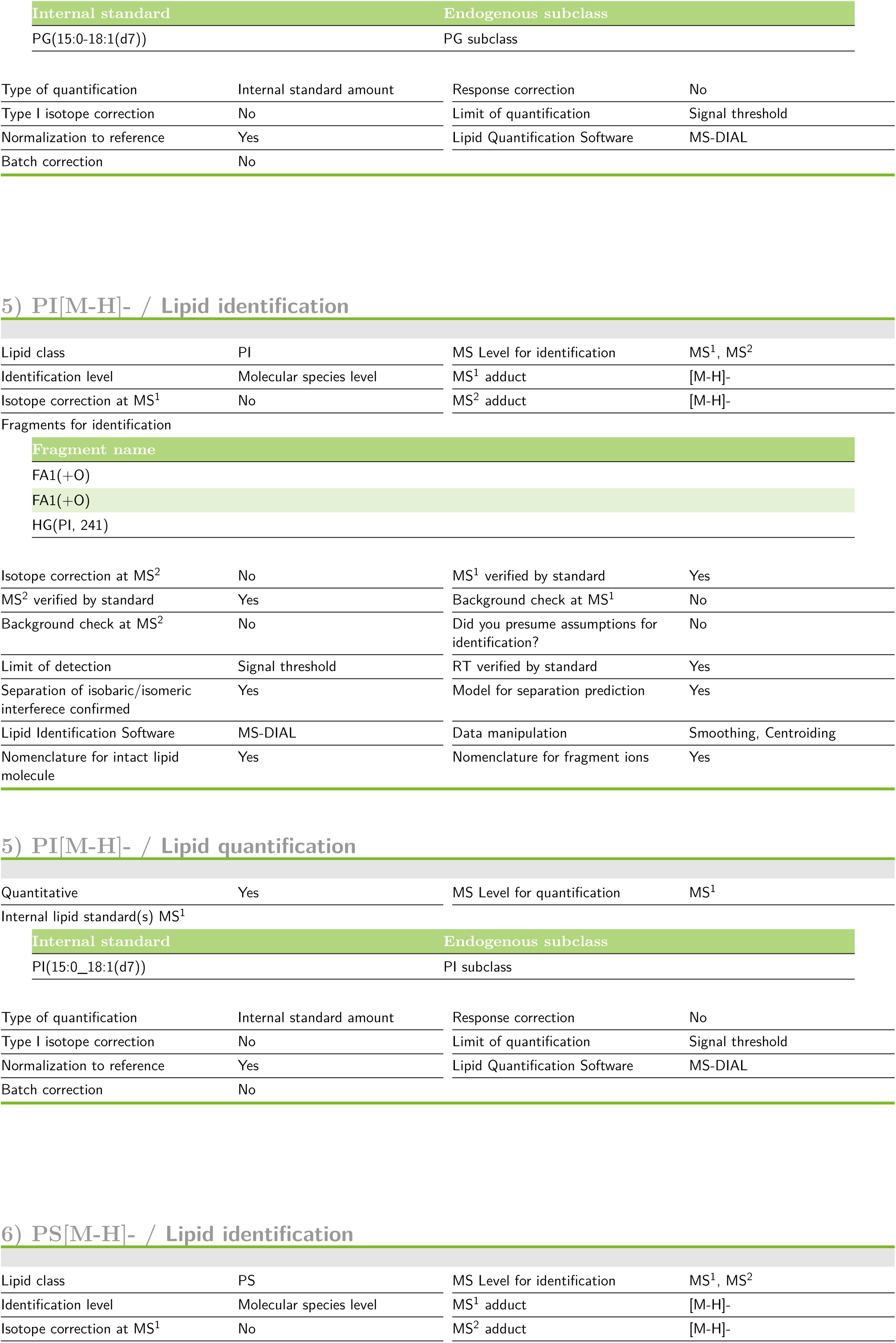
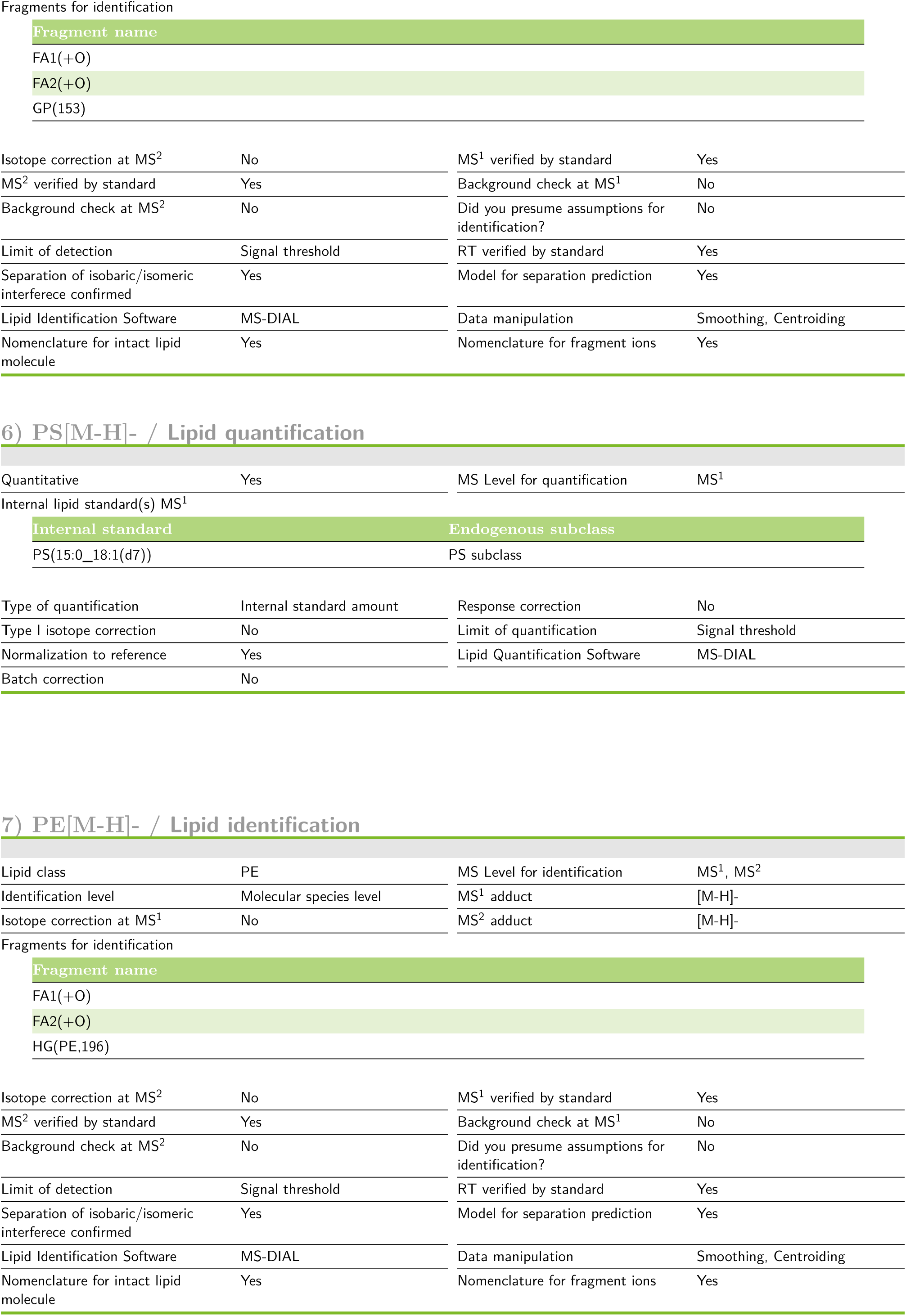
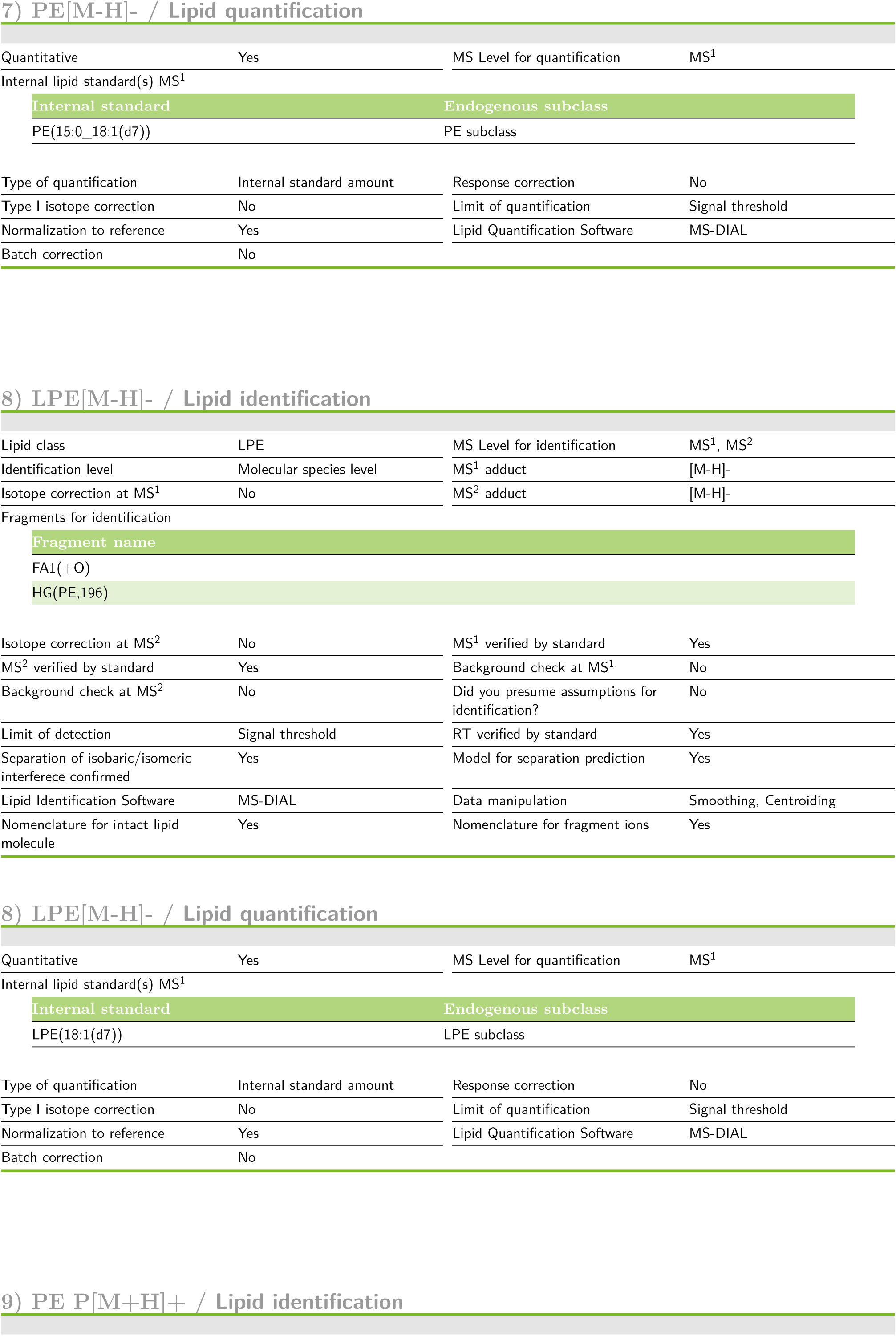
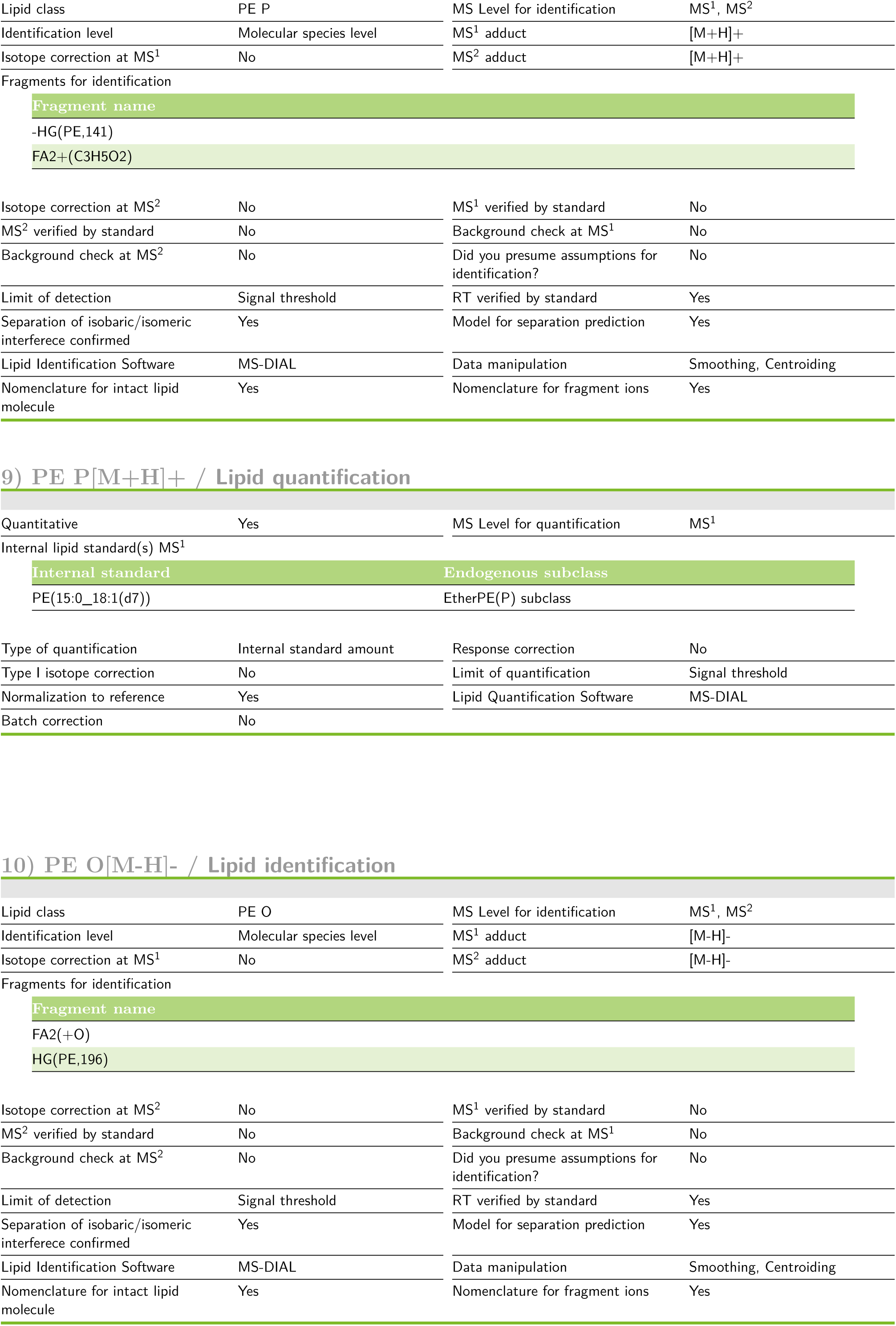
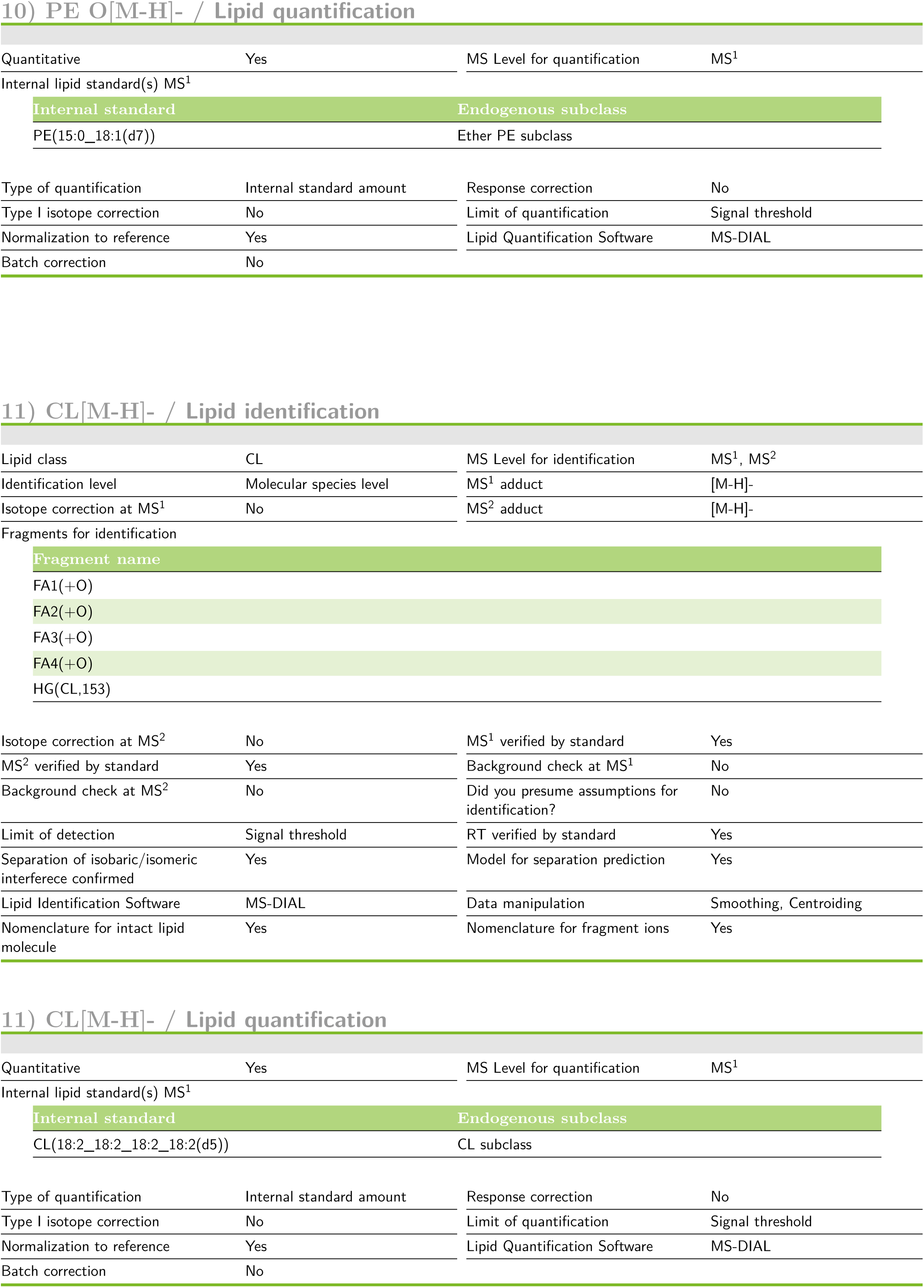
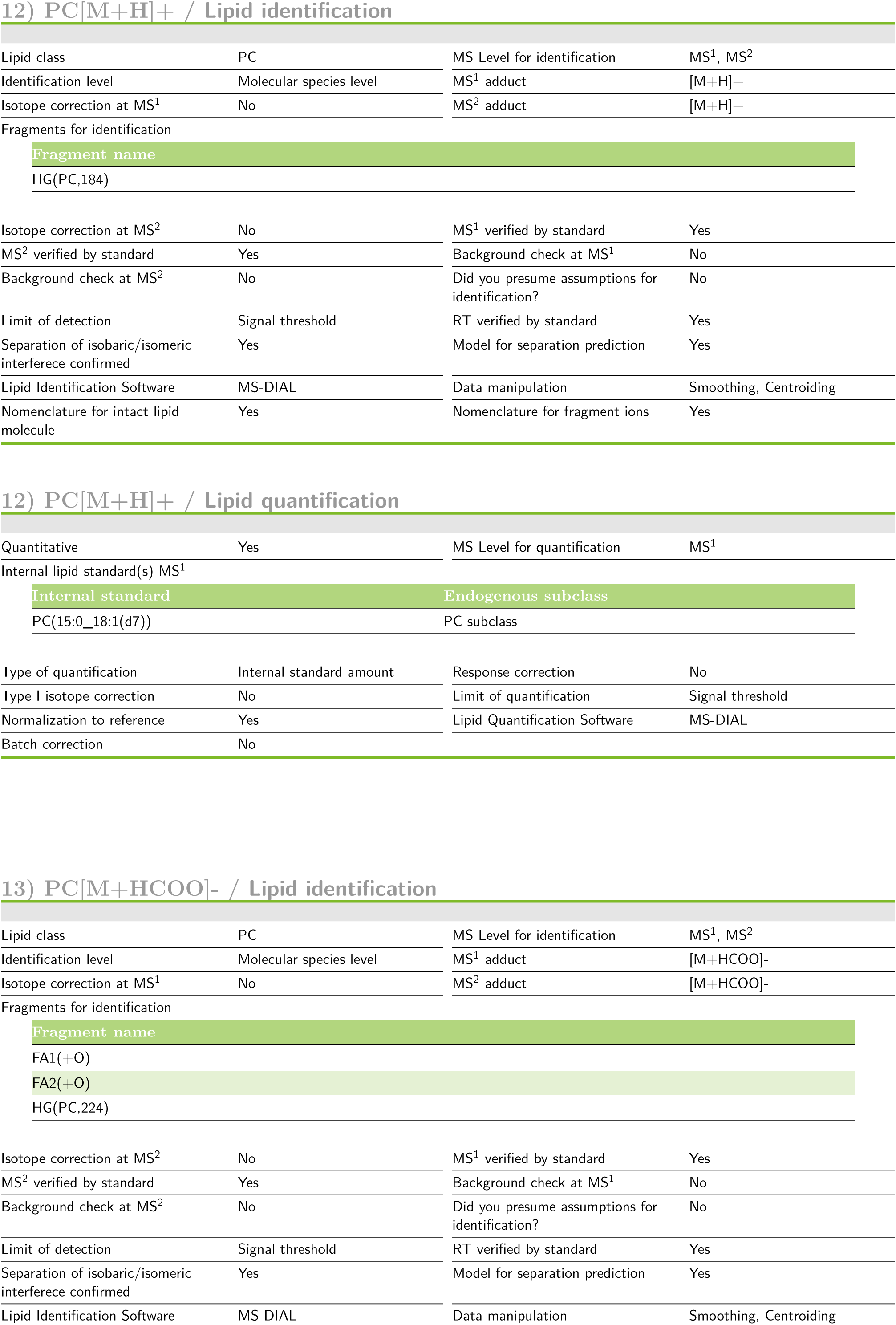
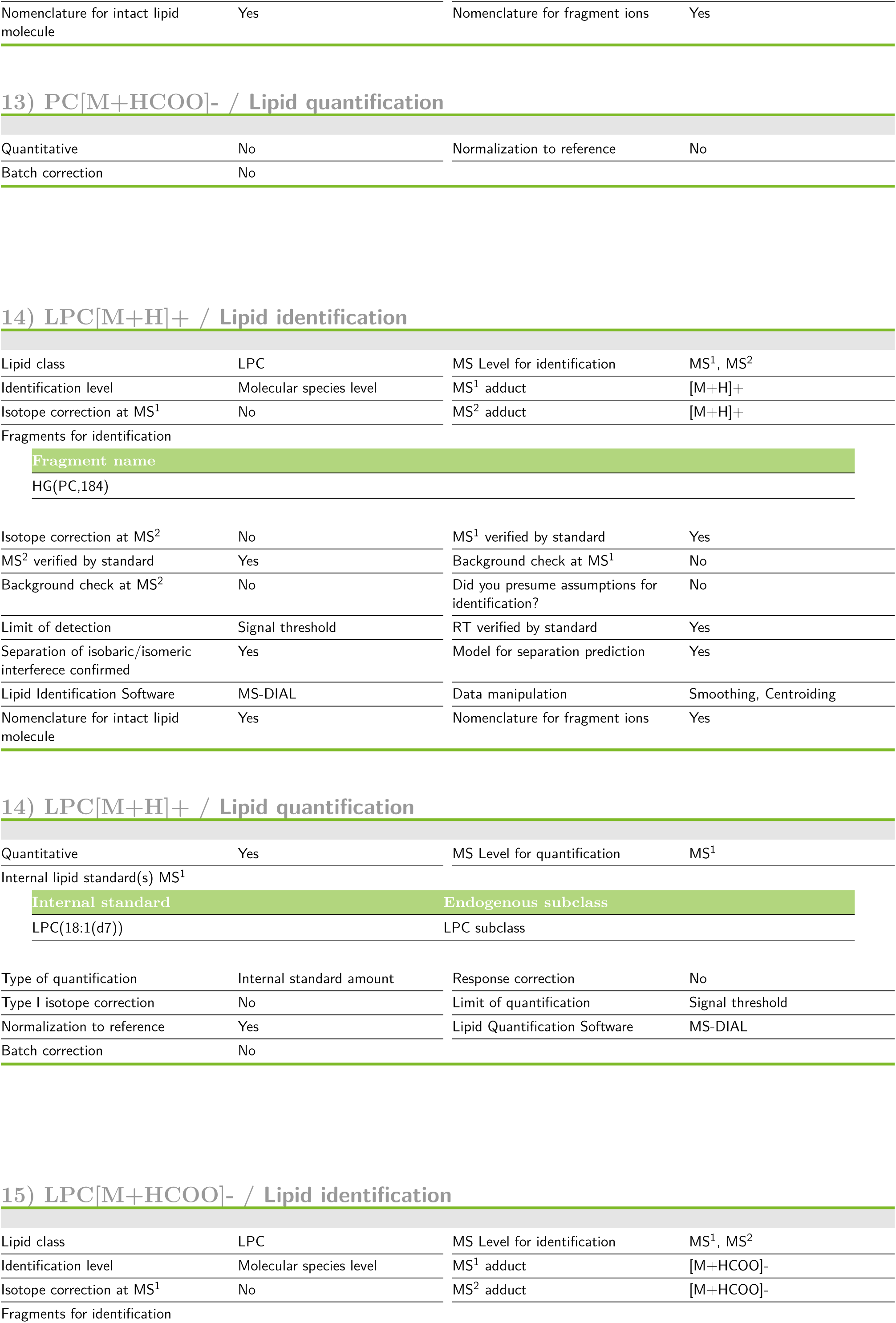
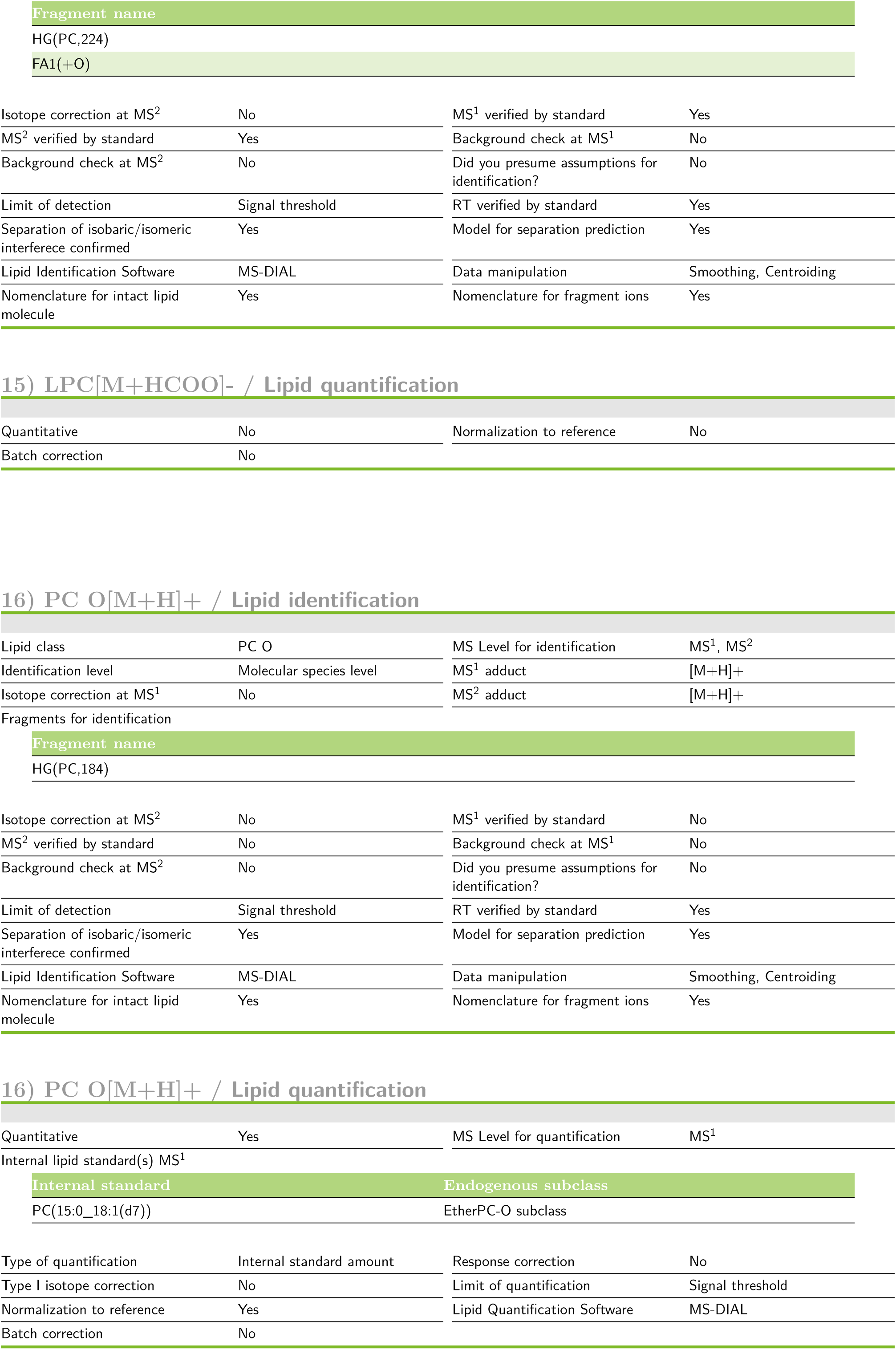
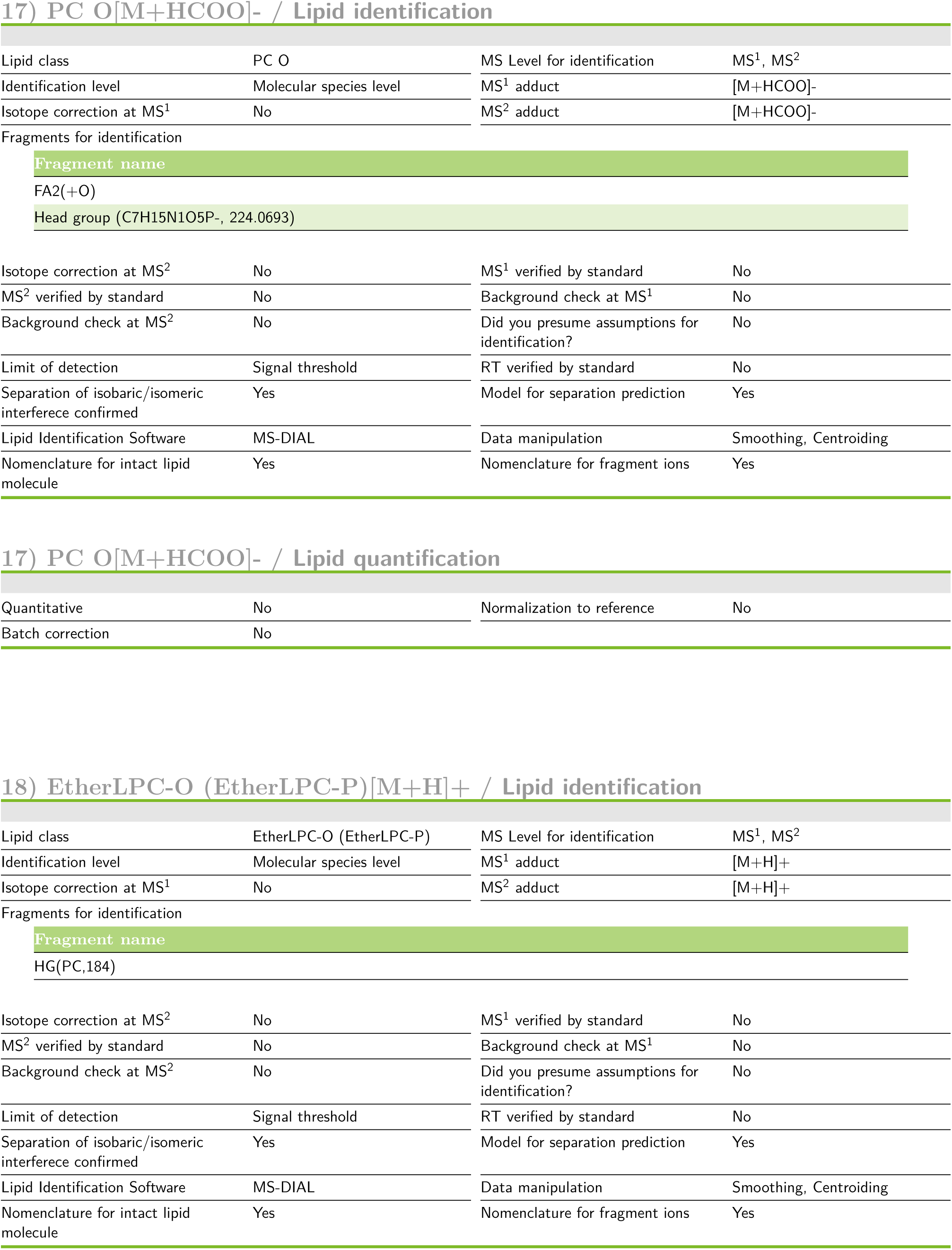
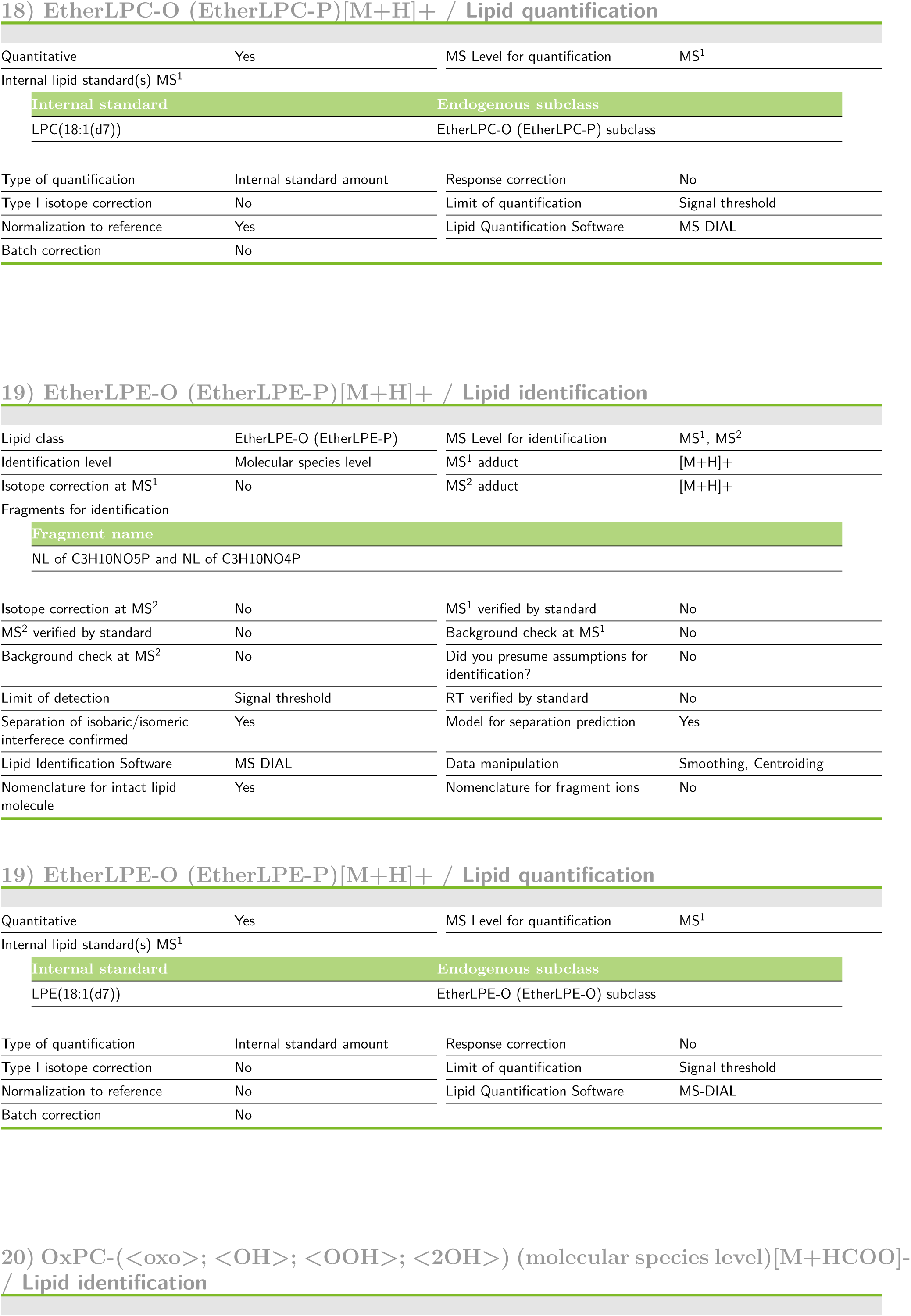
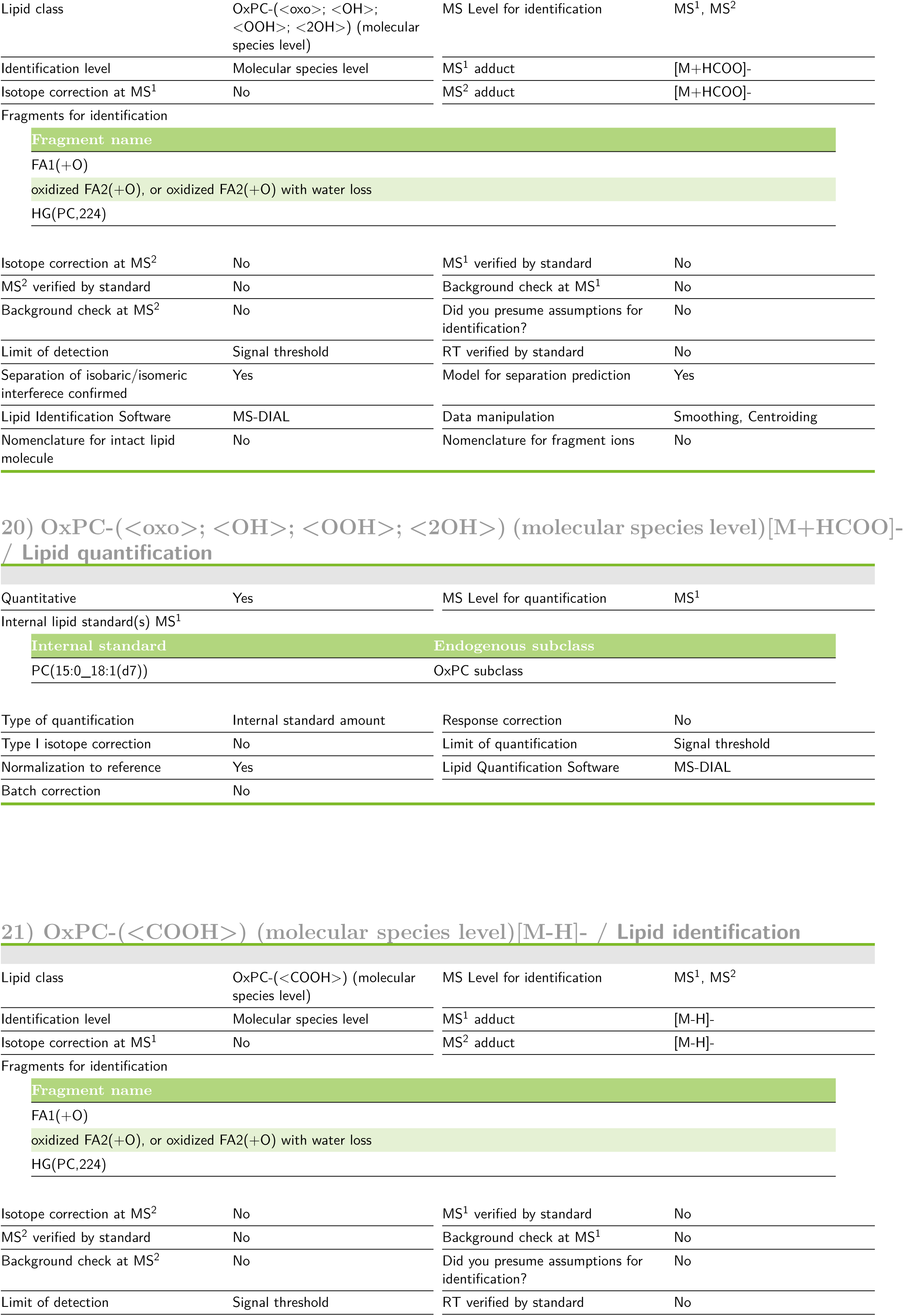
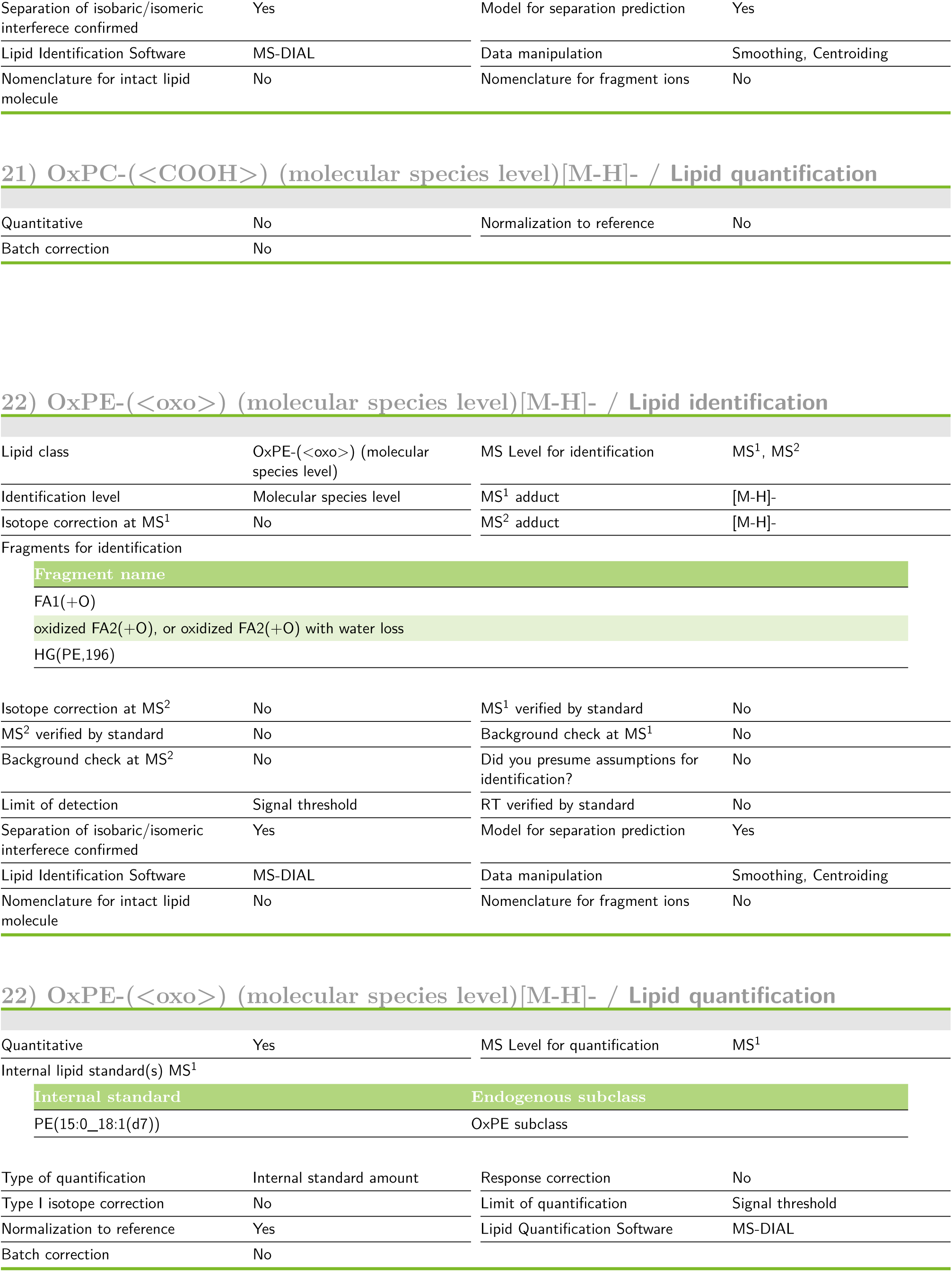
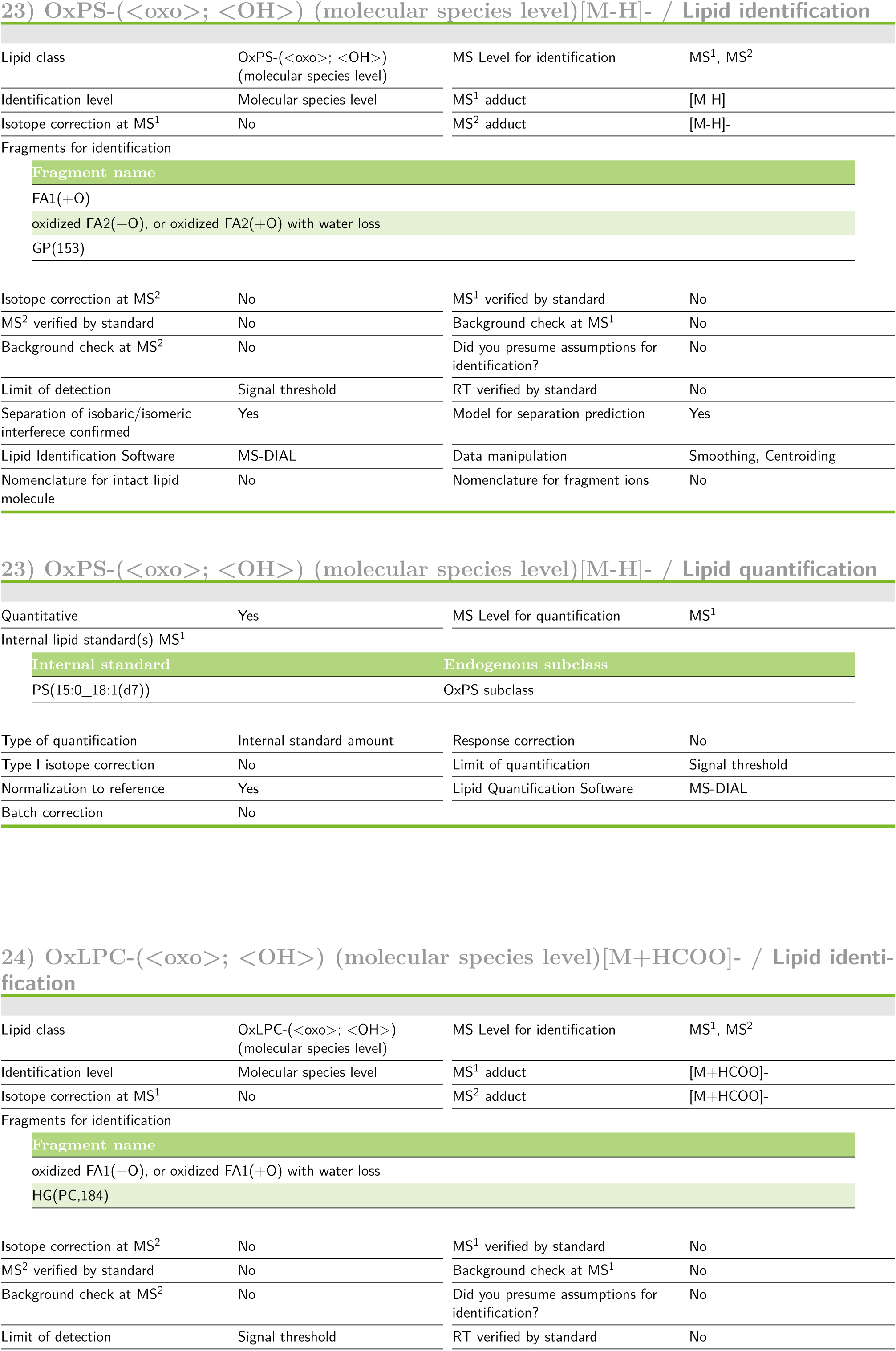
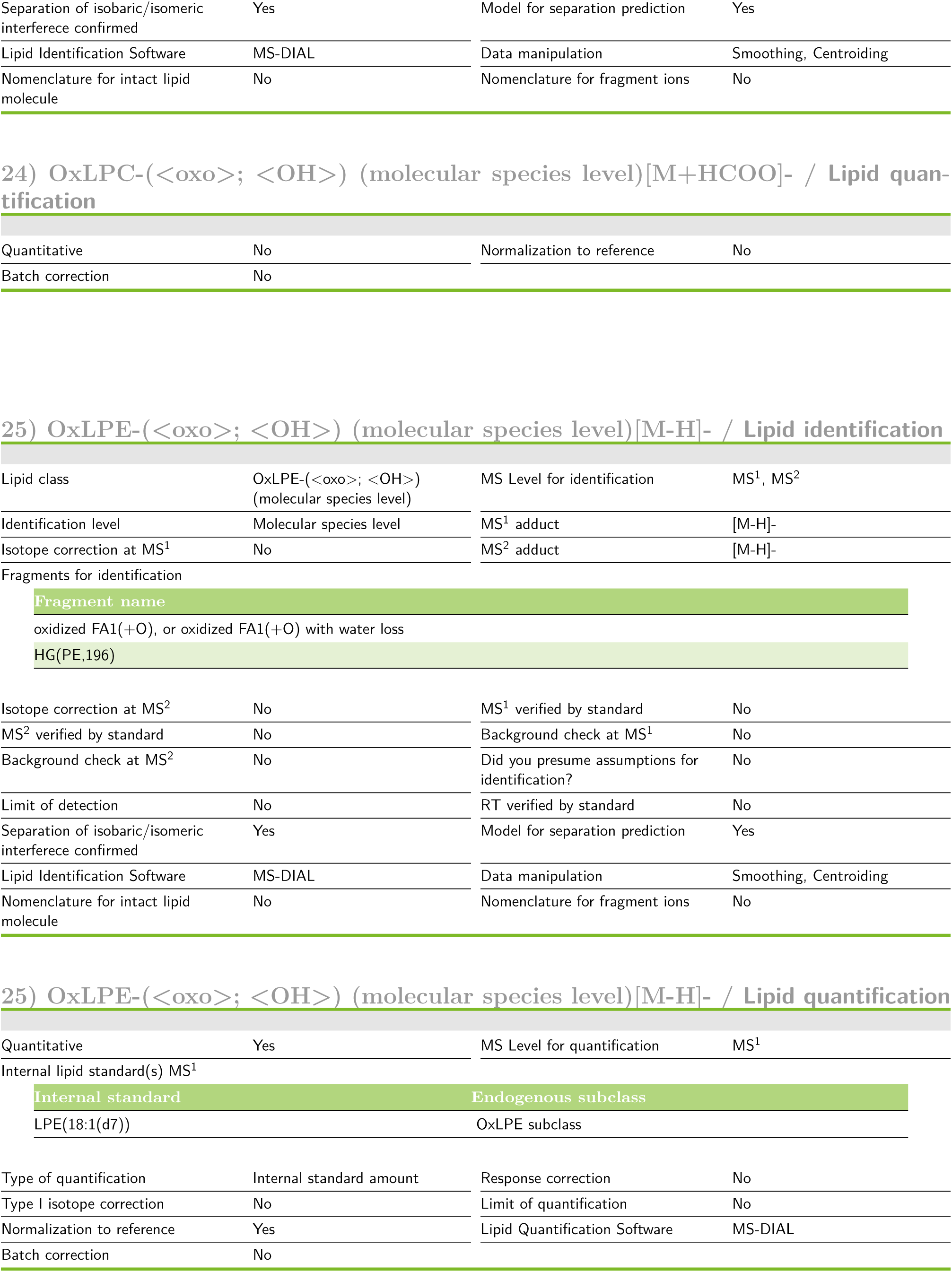
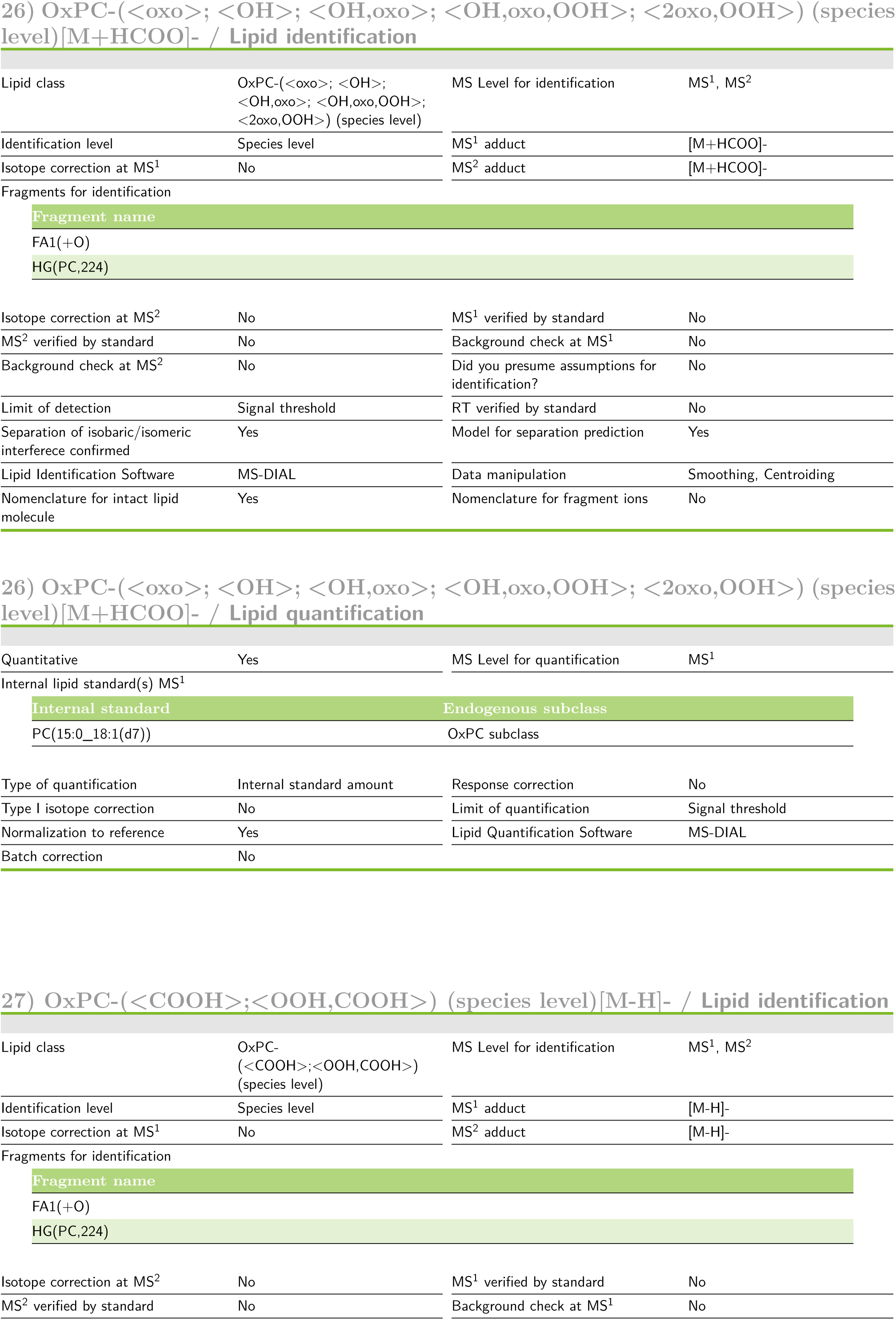
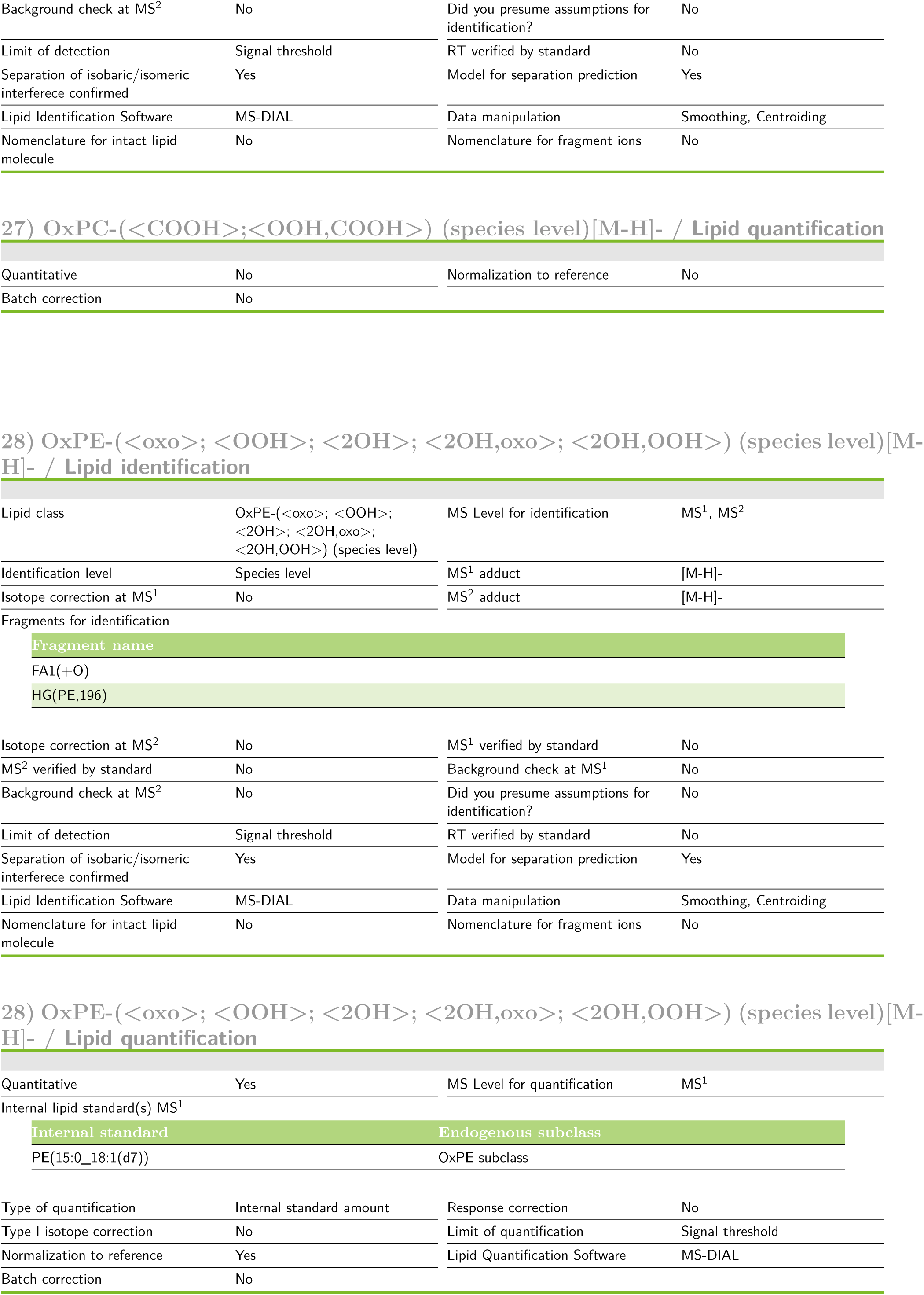
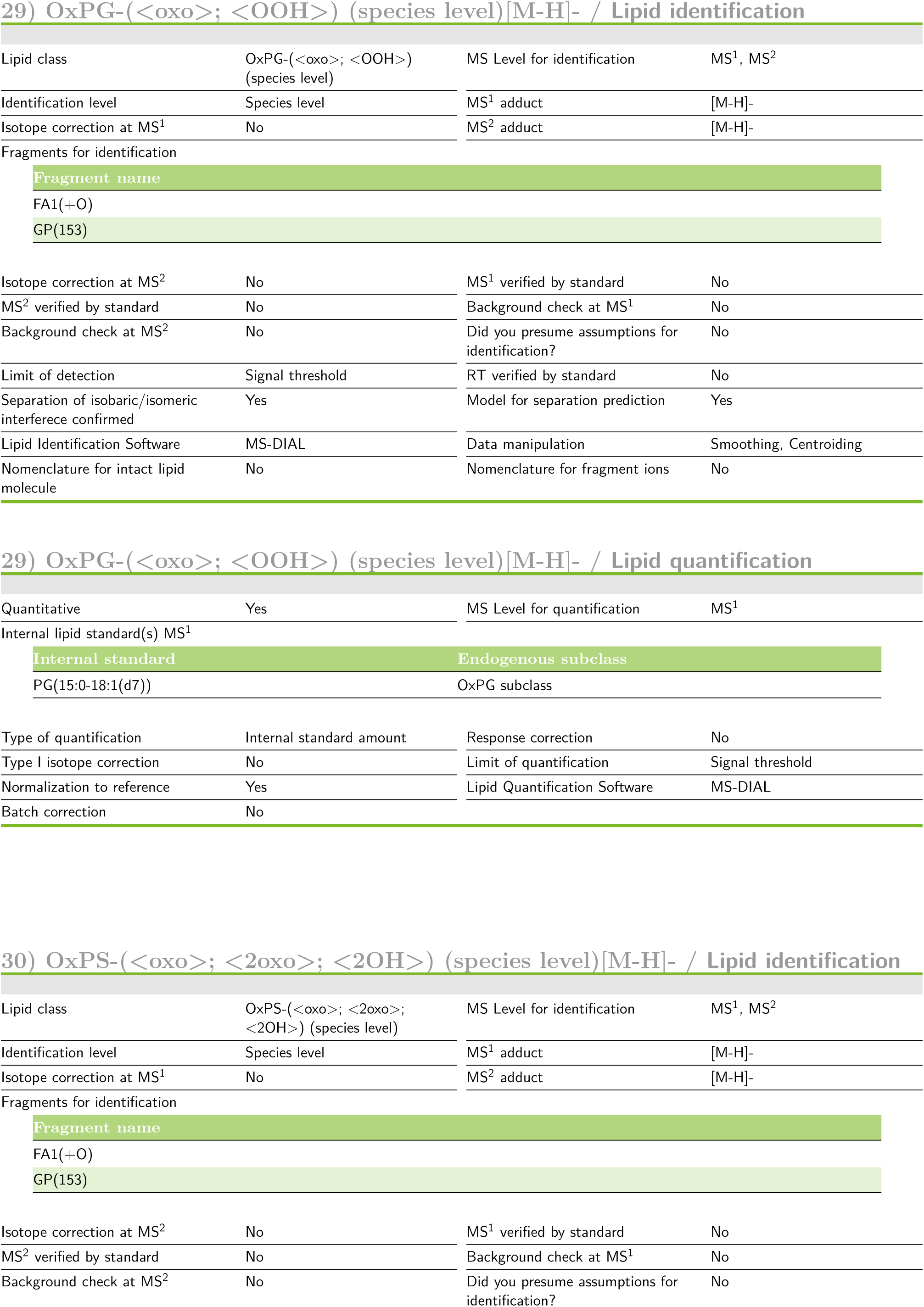
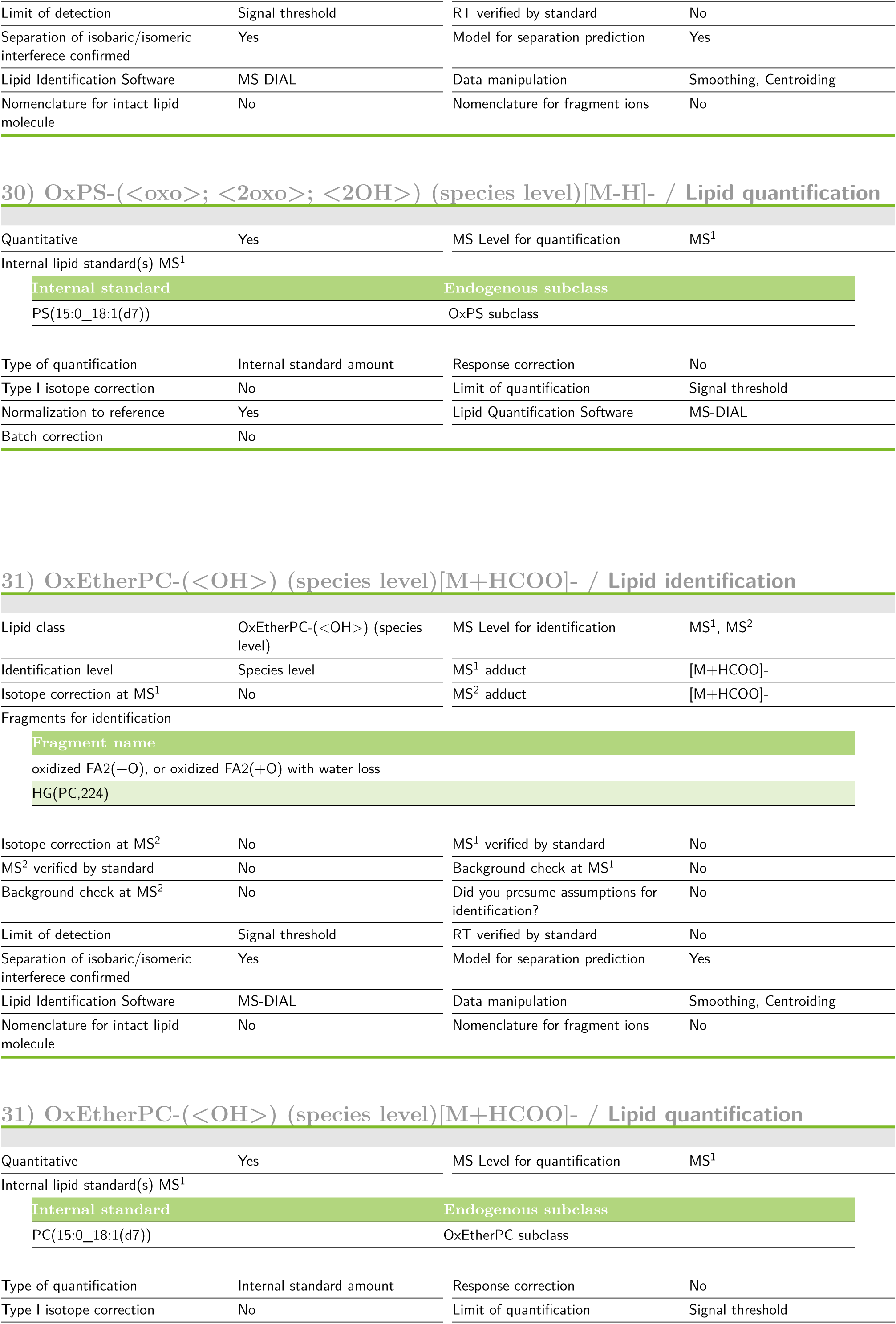
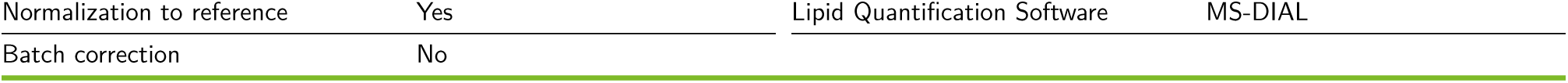
Lipidomics Standards Initiative reporting checklist.

